# Restricting visual exploration directly impedes neural activity, functional connectivity, and memory

**DOI:** 10.1101/2020.05.12.091504

**Authors:** Zhong-Xu Liu, R. Shayna Rosenbaum, Jennifer D. Ryan

## Abstract

We move our eyes to explore the visual world, extract information, and create memories. The number of gaze fixations – the stops that the eyes make – has been shown to correlate with activity in the hippocampus, a region critical for memory, and with later recognition memory. Here, we combined eyetracking with fMRI to provide direct evidence for the relationships between gaze fixations, neural activity, and memory during scene viewing. Compared to free viewing, fixating a single location reduced: 1) subsequent memory, 2) neural activity along the ventral visual stream into the hippocampus, 3) neural similarity between effects of subsequent memory and visual exploration, and 4) functional connectivity among the hippocampus, parahippocampal place area, and other cortical regions. Gaze fixations were uniquely related to hippocampal activity, even after controlling for neural effects due to subsequent memory. Individual gaze fixations may provide the basic unit of information on which memory binding processes operate.

## Introduction

The oculomotor system may be a unique effector system that supports the development of memory^1^. Key structures within the oculomotor and hippocampal memory systems are phylogenetically old structures^2, 3^, and their shared evolutionary histories has resulted in a complex network of structural and functional connections between the two systems that span temporal, parietal, and frontal regions^4^. By frequently moving the high-resolution fovea across the external world, rich visual details may be extracted, accumulated, and ultimately stored into a lasting memory representation^5–7^. Indeed, numerous eye tracking studies, across decades of research, have shown that gaze fixations – the discrete stops that are made by the eyes – predict subsequent memory, irrespective of the duration of viewing time^8–10^. This relationship appears to causal, rather than merely correlational: restricting eye movements at encoding by having participants maintain central fixation results in a decrease in subsequent recognition memory performance^10, 11^.

Findings from human neuroimaging provide further evidence of a relationship between the oculomotor and hippocampal memory systems. Neural activity in the hippocampus (HPC) increases with an increasing number of gaze fixations^12^, and this relationship is weaker in older adults who have smaller hippocampal volumes^13^. Other work shows that hippocampal activity increases with decreasing fixation duration, presumably due to a higher rate of gaze fixations, although this was not assessed directly^14^. However, the neuroimaging evidence to date is correlational in nature, as participants’ eye movements were measured under natural viewing conditions. If the relationship between fixations and hippocampally-mediated memory is direct, then modifying rate and extent of visual exploration should modulate activity in the HPC as well as subsequent memory. Likewise, given the vast structural and functional network within which the oculomotor and hippocampal memory systems interact^4, 15^, changes in patterns of visual exploration should also affect either the set of regions that comprise functionally connected networks, or the strength of those functional connections.

In the present study, we experimentally manipulated participants’ viewing behavior and observed changes in the functional engagement of the HPC, surrounding medial temporal lobe, and ventral visual stream. Participants studied scenes and scrambled images either under *free-viewing* conditions, or they were asked to maintain fixation during viewing (*fixed*-*viewing*). Following scanning, participants were given a recognition memory task. Based on prior work, it was expected that the number of gaze fixations would relate to subsequent recognition memory, and that *fixed-viewing* would result in a decrease in recognition memory^10, 11^. We also predicted that gaze fixations would be predictive of neural responses in the HPC, and in the parahippocampal place area (PPA) due to the use of scenes in the present task^16–18^. In comparison to *fixed-viewing, free-viewing* was predicted to result in increases in neural responses along the ventral visual stream and into the medial temporal lobe and HPC. Manipulating visual exploration was further predicted to change the configuration of functional networks and/or the degree to which regions were functionally connected during task performance. Finally, conjunction and cross-voxel similarity analyses were conducted to determine the extent to which the influence of gaze fixations on neural activity was similar to, or was unique from, activity indicative of successful encoding.

Alterations in the relationships between gaze fixations, neural activity in the hippocampus and broader medial temporal lobe, and subsequent memory due to viewing manipulations would suggest that visual exploration is a general mechanism that supports the binding of information into memory. Further evidence for the integral link between visual exploration and hippocampal function are expected to be found in patterns of brain activity that are similar across gaze modulation and subsequent memory effects. Unique effects of gaze fixations on hippocampal activity, above and beyond effects of subsequent memory, may suggest that gaze fixations provide the basic units of information on which the hippocampus operates.

## Results

### Eye Movements

To ensure that participants had complied with the viewing instructions (Figure 1), we compared the number of fixations (Figure 2A, 2B, and 2C) and the average saccade amplitude (Figure 2D) in the *free-* and *fixed-viewing* conditions for the scenes and scrambled pictures. A 2×2 repeated measures ANOVA (viewing condition by stimulus type) revealed that participants made a greater number of gaze fixations (*F*(1,35) = 97.24, *p* < .0001), and had a larger saccade amplitude (*F*(1,35) = 80.53, *p* < .0001), in the *free-viewing* condition than the *fixed-viewing* condition, for both the scenes and the scrambled pictures. A significant interaction for the number of fixations only (*F*(1,35) = 8.78, *p* = .005) indicated that more gaze fixations were made to scenes versus scrambled pictures in the *free-viewing*, but not in the *fixed-viewing* condition.

**Figure 1.**
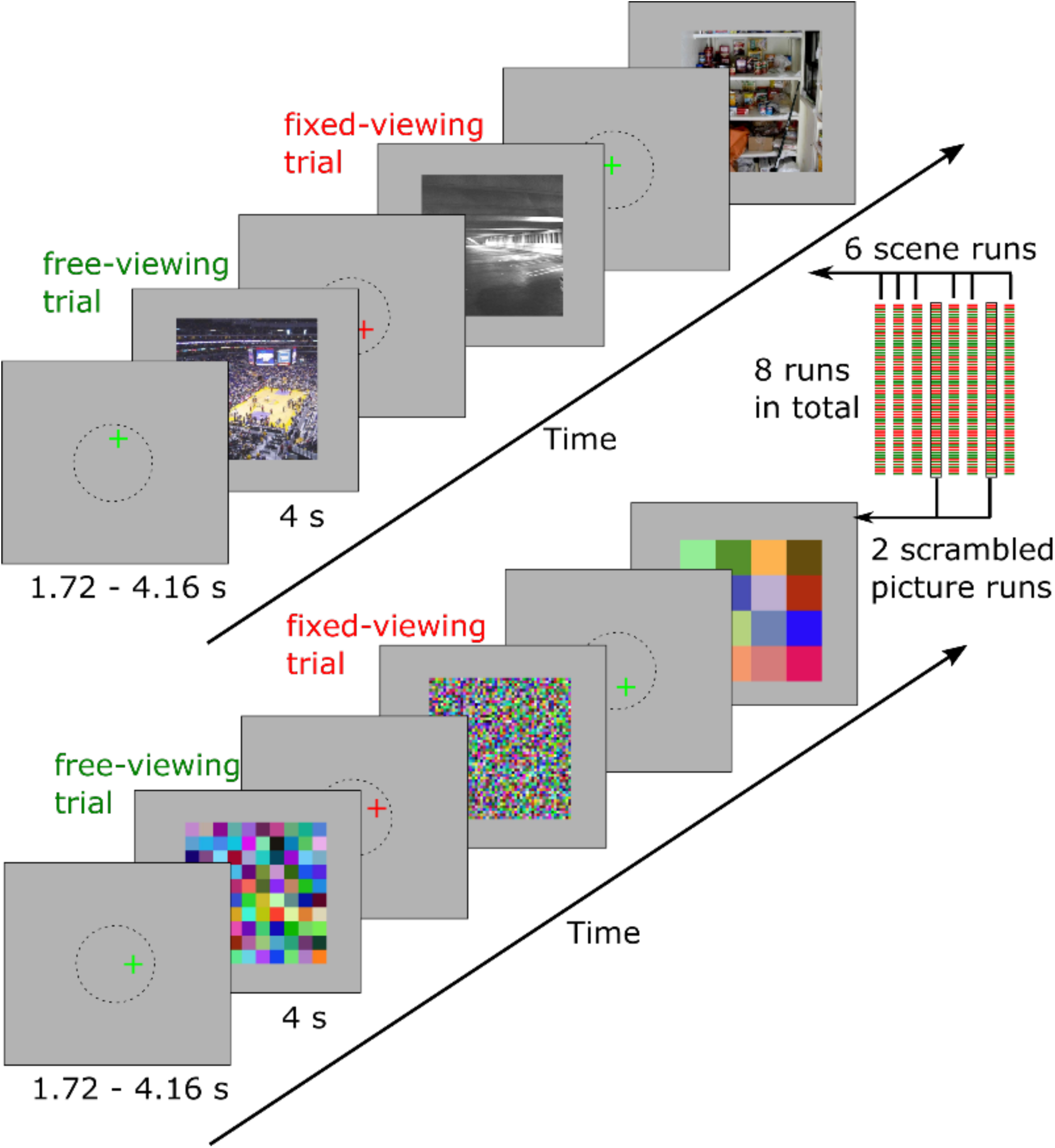
Participants were presented with either a green or red fixation cross at the start of each encoding trial. A green fixation cross instructed the participants to freely view the upcoming scene or scrambled image (*free-viewing)*. A red fixation cross instructed the participants to maintain fixation on the location of the cross during the presentation of the upcoming scene or scrambled image (*fixed-viewing*). Participants were instructed to encode as much information regarding the scene or scrambled image as possible under both viewing conditions. Six runs of scenes and two runs of scrambled images were presented to the participants, with timing as indicated.

**Figure 2.**
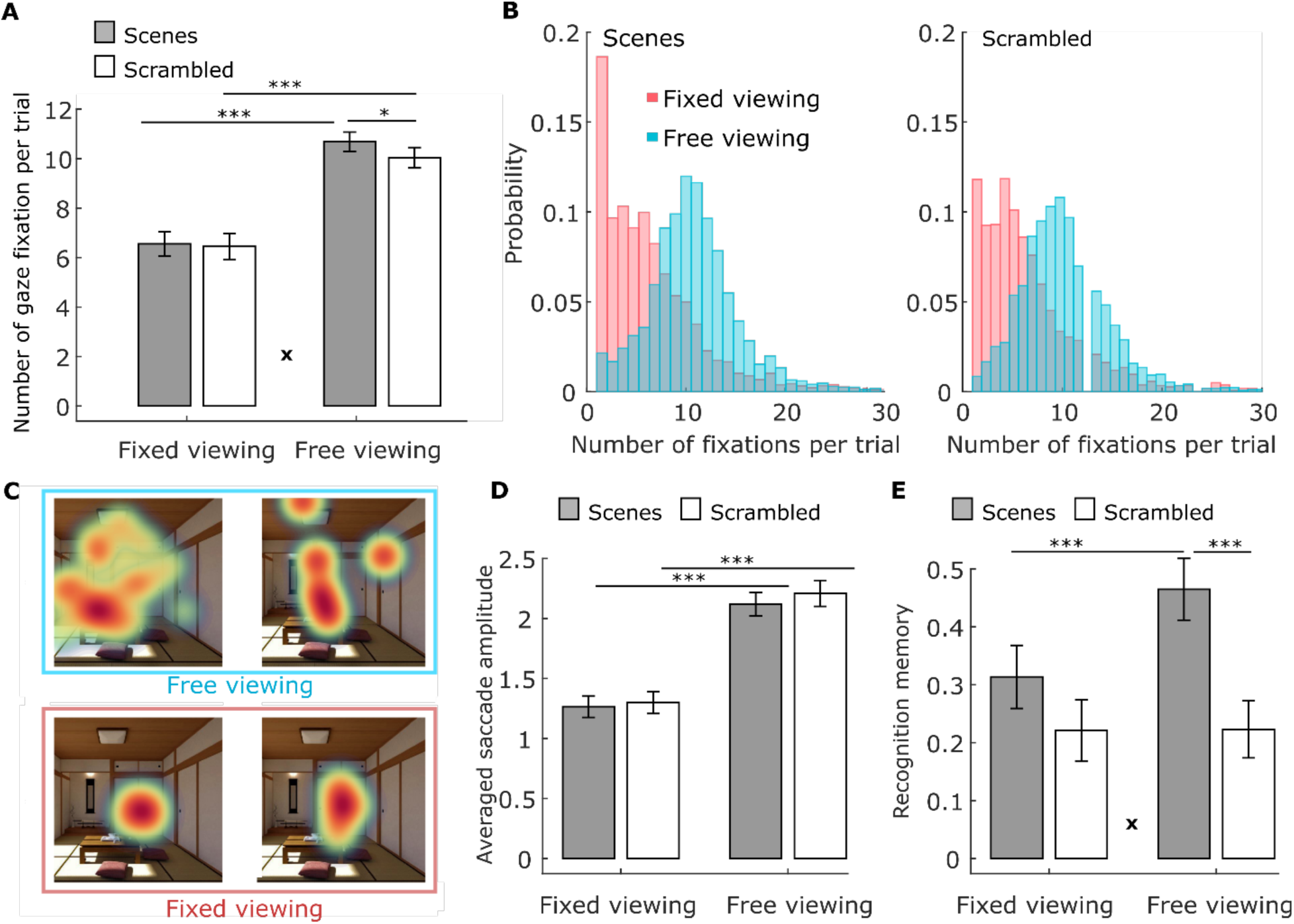
**A**. Number of gaze fixations. Participants made more fixations on average per trial under *free-viewing* versus *fixed-viewing* instructions. More fixations were elicited by scenes versus scrambled pictures, but only in the *free-viewing* condition. **B**. The distribution (across all participants) of the number of gaze fixations for the scenes (left) and scrambled pictures (right) under fixed (red) and free (blue) viewing conditions. Similar distributions are observed for the number of fixations for the scenes and scrambled pictures. **C**. Fixation heatmaps (weighted by fixation durations) for a sample scene in the *free-viewing* versus *fixed-viewing* condition illustrates the difference in viewing patterns in the two conditions. **D**. Average saccade amplitude, which is larger under *free-viewing* versus *fixed-viewing* instructions. **E**. Recognition memory performance (with confidence considered) was better in the *free-viewing* vs. *fixed-viewing* condition only for scenes. * = *p* < 0.05; ***= *p* < 0.001. **x** = significant ANOVA interaction effect, *p* < .005.

### Memory Performance

A 2 × 2 repeated measures ANOVA on recognition memory performance (with confidence considered) revealed a significant interaction (*p* < .05), indicating that recognition memory was significantly better in the *free-* versus *fixed-viewing* condition only for the scenes (Figure 2E). Similar results were obtained when memory was measured using hit rate – false alarm rate. Detailed statistics see Supplementary Table S1). We then ran a linear regression analysis for each condition in each run using the trial-wise number of fixations to predict recognition memory (with confidence considered). The number of gaze fixations positively predicted recognition memory for the scenes (*t* = 2.75, *p* = .009 and *t* = 6.10, *p* <.0001 for the *fixed-* and *free-viewing* condition, respectively), but not for the scrambled pictures (*p* > .5). The prediction was stronger for scenes studied under *free-* versus *fixed-viewing* instructions (*p* = .0013. See Supplementary Table S1).

### Neuroimaging results

#### Brain activation differences between the free-vs. fixed-viewing conditions

To assess whether activity in the PPA and HPC was modulated by viewing instructions, we examined the brain activation contrast between the *free-* and *fixed-viewing* conditions for the scenes and scrambled pictures. The PPA and HPC showed stronger activation in the *free-viewing* condition than in the *fixed-viewing* condition for scenes (Figure 3A; *t* = 5.20/4.67 and 9.63/10.32, for the left/right HPC and left/right PPA respectively, all *p* <.0001). Similar effects were found for the viewing of scrambled pictures (Figure 3A; *t* = 4.61/3.64 and 7.16/8.17, for the left/right HPC and left/right PPA respectively, all *p* <.001). Directly comparing the effect of viewing condition for the scenes and scrambled pictures revealed that the *free*-versus *fixed-viewing* difference in activation for the left and right PPA was larger for scenes than scrambled pictures (*t* = 6.25 and 6.79, *p* <.0001). When the whole HPC was used as the ROI, the effect of viewing condition did not differ between scenes and scrambled pictures (*t* = .14 and .17, *p* = .80 and .51, for the left and right HPC, respectively).

**Figure 3.**
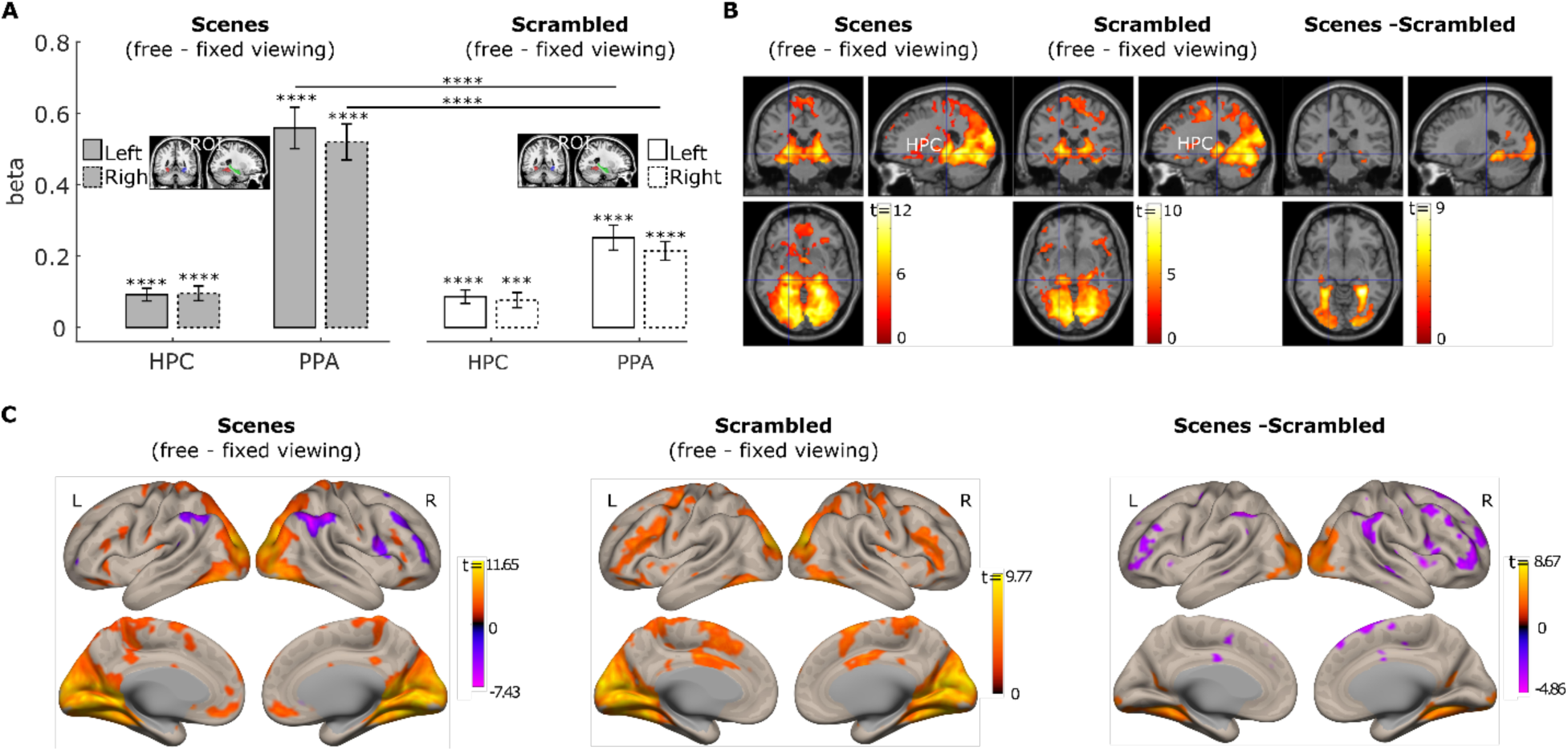
**A**. Neural activity difference between the *free-* and *fixed-*viewing in the hippocampus (HPC) and parahippocampal place area (PPA) ROIs. Significant responses were observed in the HPC and PPA bilaterally for the *free*-versus *fixed*-viewing contrast. This *free-* versus *fixed* viewing difference was larger for scenes than for scrambled pictures in the PPA, but not in the HPC at the ROI level. ***= *p*<.001; ****=*p*<.0001. **B**. Brain section views of the *free-* versus *fixed* viewing activation differences for scenes (left), and scrambled pictures (middle). The scene (free-fixed) versus scrambled pictures (free-fixed) contrast is depicted on the right. **C**. Surface views of the *free-* versus *fixed* viewing activation differences for scenes (left), and scrambled pictures (middle). The scene (free-fixed) versus scrambled pictures (free-fixed) contrast is depicted on the right. The occipital cortex and ventral and medial temporal cortex showed stronger activation during the *free-viewing* condition, compared to the *fixed-viewing* condition. Medial temporal lobe regions and the PPA showed a stronger effect of viewing condition for the scenes compared to the scrambled pictures. For illustration purposes, data in B and C are thresholded at *p* = .005, 10-voxel extension with no corrections.

However, we also examined whether any voxels inside the HPC showed differences in the effect of viewing condition for scenes versus scrambled pictures. Based on the whole brain voxel-wise results (*p* < .005 with 10 voxel extension) and the small-volume-correction analysis in SPM 12, we found that voxels in both the left and right MNI HPC masks showed a stronger viewing condition effect (*free-viewing* > *fixed-viewing*) for scenes compared to the scrambled pictures (*p*_fwe-corr_ = .022 with family-wise error correction, *p* < .0001 without correction, peak voxel location: [−34, −32, −12] for the left HPC, and *p*_fwe-corr_ = .024 with family-wise error correction, *p* < .0001 without correction, peak voxel location: [−34, −34 – 10], for the left HPC). Brain activation section and surface images (with *p* = .005, 10 voxel extension; no corrections) in Figures 3B and 3C, respectively, illustrate the effects in these ROIs and other brain regions. The whole brain voxel-wise results are also presented in Supplementary Table S2, S3, and S4.

#### Modulation of the number of gaze fixations on neural responses

To replicate our prior work ^12, 13^, we investigated whether the trial-wise number of fixations was associated with HPC and PPA activation during *free-viewing* of scenes. Our parametric modulation ROI analysis showed that greater numbers of gaze fixations predicted stronger activation in the left and right PPA (*t* = 5.31 and 6.64, *p* <.001). The effect was not significant for the HPC when the whole HPC anatomical ROIs were used (*t* = .68 and 1.12, *p* =.50 and .27 for the left and right HPC respectively; Figure 4A). However, not all HPC regions along its longitudinal axis were affected by the viewing manipulation (Figure 3B). Therefore, based on the whole brain voxel-wise results (*p* < .005 with 10 voxel extension), we did a similar small volume correction analysis using the HPC MNI template masks to correct for multiple comparisons and found a cluster of voxels in the left and right HPC with peak voxels that survived family-wise error correction at *p*_fwe-corr_ = .05 (Figure 4B). The whole brain voxel-wise results are presented in Figure 4C and Supplementary Table S5. Fixation modulation effects were not significant in HPC and PPA ROIs in the scene *fixed-viewing*, scrambled *free-viewing*, and scrambled *fixed-viewing* conditions (Figure 4D, 4E, and 4F).

**Figure 4.**
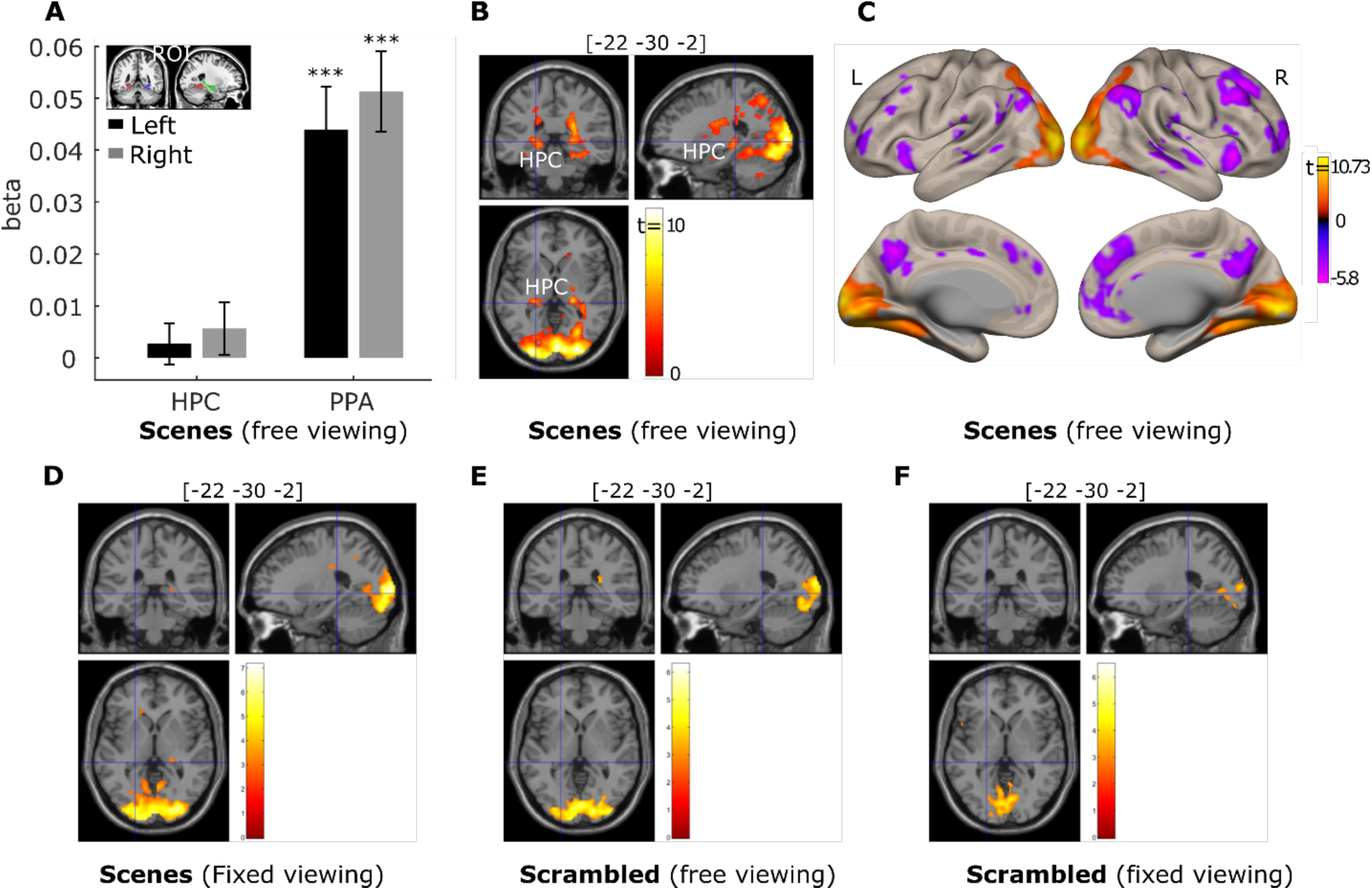
The number of gaze fixations positively predicts activation in the hippocampus (HPC) and parahippocampal place area (PPA) bilaterally. **A**. Results for the anatomical HPC ROIs and functional PPA ROIs (***= *p*<.001, two-tailed t test). **B**. Activation of a cluster in the bilateral HPC was positively predicted by the number of gaze fixations (*p*<.05 Small volume family-wise error correction). For the left HPC: *p*_fwe-corr_ = .001, *p* < .00001, voxel size = 177 voxels with the peak voxel location = [−22, −30, −2]; for the right HPC: *p*_fwe-corr_ = .009, *p* < .00001, voxel size = 87 voxels with the peak voxel location = [28, −34, 4]. **C**. Brain surface views for the prediction of gaze fixations. **D, E**, and **F** show the fixation modulation effects for the scenes under *fixed viewing*, scrambled pictures under *free viewing*, and scrambled pictures under *fixed viewing*, respectively. For illustration purposes, brain section (**B, D, E, F**) and surface (**C**) views are also presented at *p*<.005, 10-voxel extension with no corrections. For the brain surface views, L indicates left hemisphere and R the right hemisphere.

#### PPA connectivity differences due to viewing condition

Given its role in the perceptual processing of scenes, we used the bilateral PPA as seeds to explore the effect of viewing condition on functional connectivity. Using the generalized psychophysiological interaction (gPPI) analysis implemented in the CONN toolbox, we found voxel clusters in the bilateral HPCs whose connectivity with the PPA was stronger in the *free-viewing* than *fixed-viewing* condition (Threshold: Small volume correction, *p*_fwe-corr_ < .05. Figure 5). Other regions, such as the lateral and medial occipital cortex and inferior and superior parietal lobules, also showed stronger connectivity with the PPAs in the *free-* versus *fixed-viewing* condition. The whole brain voxel-wise PPA connectivity results are also presented in Supplementary Table S6.

**Figure 5.**
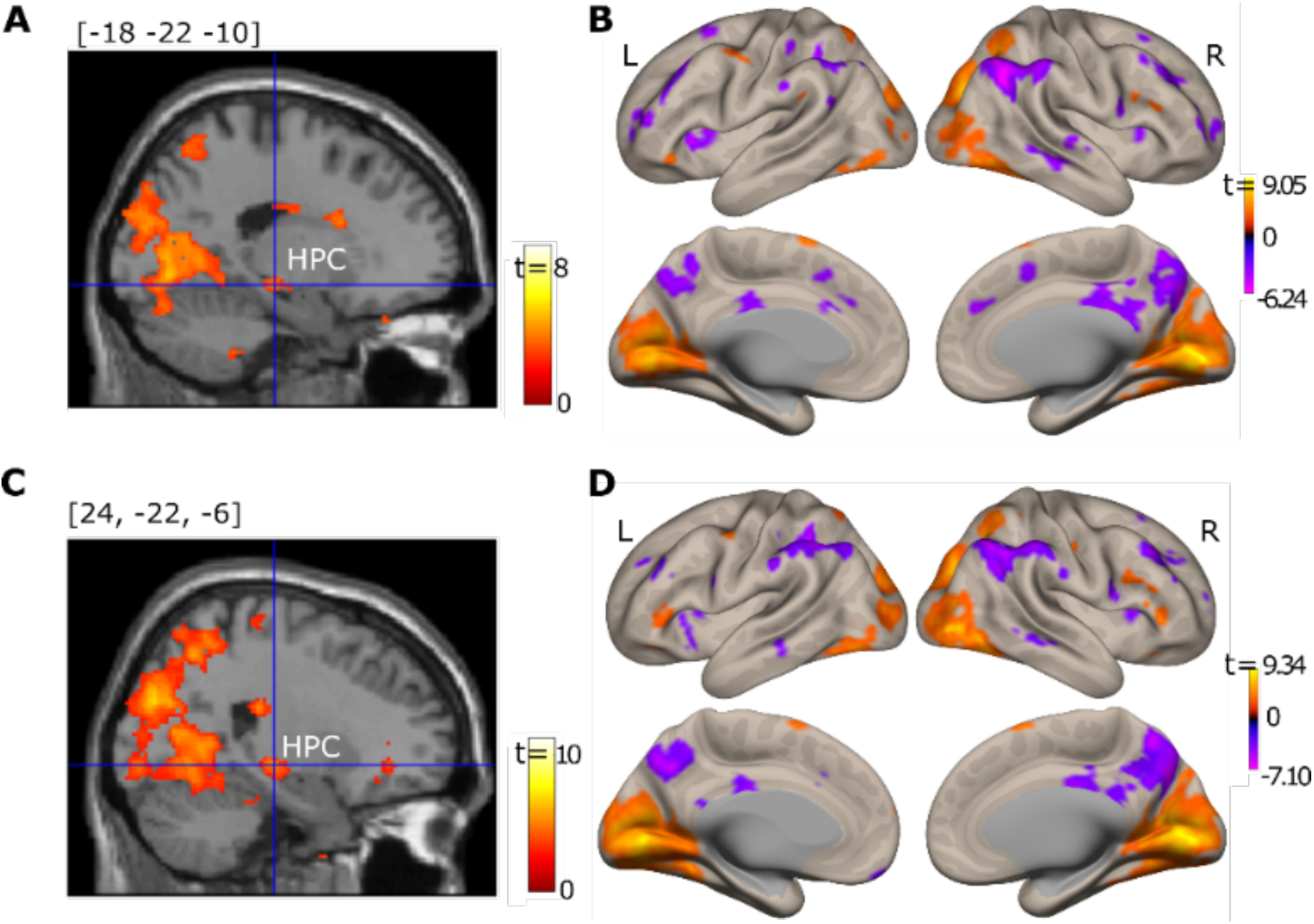
Brain section views of the left (**A**) and right (**C**) HPC, for which connectivity with the left (top) and right (bottom) PPA was modulated by viewing condition. Brain surface viewings of free versus fixed viewing modulation effect on functional connectivity for the left (**B**) and right (**D**) PPA. For illustration purposes, data were thresholded at *p* = .005, 10-voxel extension with no corrections. The left PPA showed connectivity with a cluster in the left HPC (voxel size = 20 voxels with the peak voxel location = [−22, −22, −10], *p*_fwe-corr_ = .016, *p* < .001), and the right PPA showed connectivity with a cluster in both the left HPC (voxel size = 58 voxels with the peak voxel location = [−22, −24, −8], *p*_fwe-corr_ = .001, *p* < .0001) and the right HPC (voxel size = 24 voxels with the peak voxel location = [26, −24, −8], *p*_fwe-corr_ = .014, *p* < .001) that was stronger in the *free-* than in the *fixed-viewing* condition.

#### Modulation of the number of gaze fixations and subsequent memory

The relationship between the modulation effects of the number of gaze fixations and of subsequent memory performance in the *free-viewing* scene condition was examined. First, using parametrical modulation ROI analysis, we found that PPA activation was stronger for subsequently remembered than forgotten trials (*t* = 5.17 and 5.81, *p* <.001, one-tailed; Figure 6A). The right HPC showed similar significant effects (*t* = 1.75, *p* = .045, one-tailed; left HC, *t =* 1.36, *p* =.092, one-tailed; Figure 6A). The whole brain voxel-wise subsequent memory effects are presented in Supplementary Table S7.

**Figure 6.**
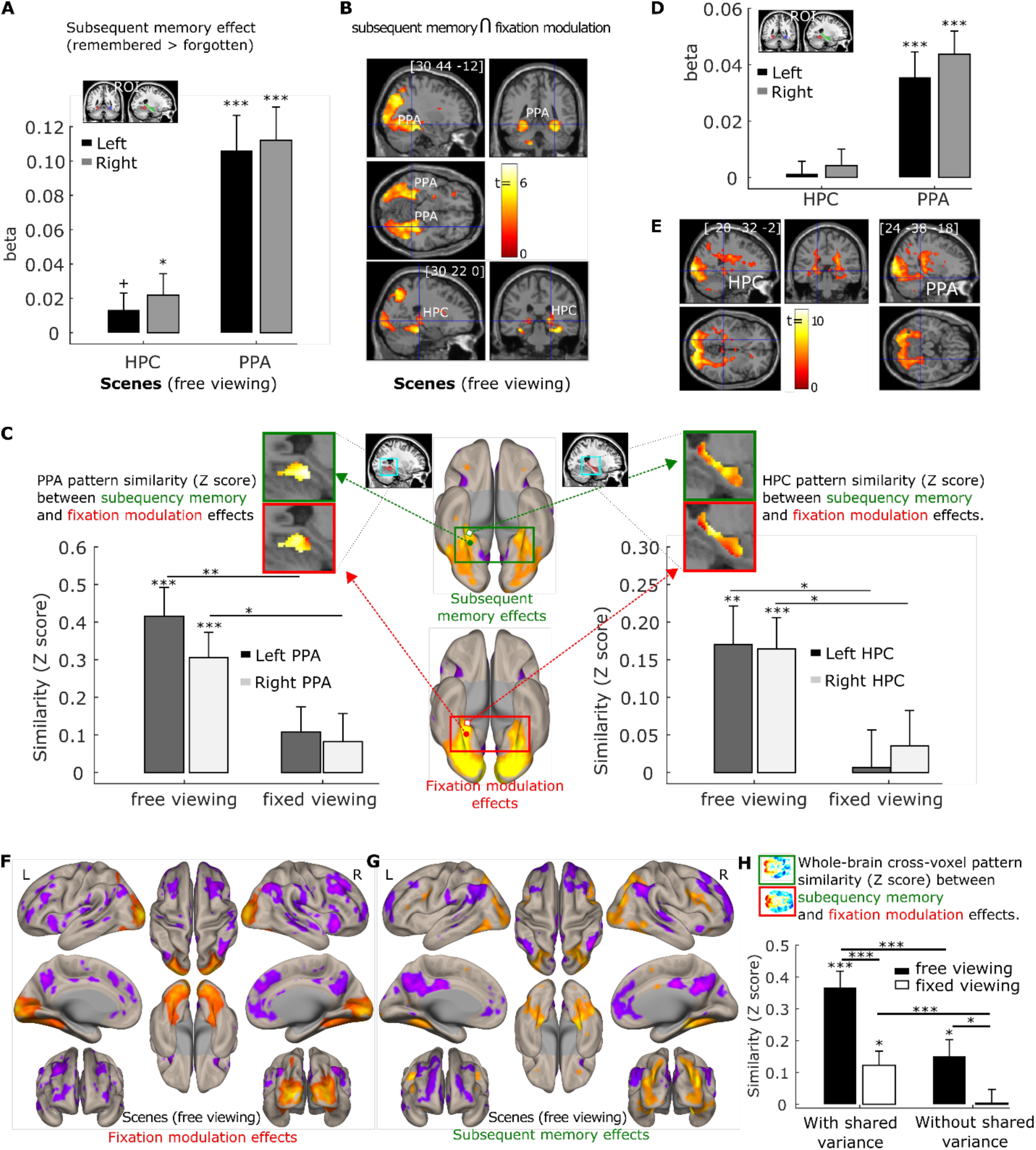
**A**. Subsequent memory positively predicts activation in the hippocampus (HPC) and parahippocampal place area (PPA) bilaterally. ROI analysis results were based on anatomical HPC ROIs and functional PPA ROIs. **B**. Activation in a cluster in the PPA bilaterally and in the right HPC was positively predicted by the number of gaze fixations and subsequent memory in a conjunction analysis (*p*=.0025, with 50-voxel extension, no corrections). **C**. Cross-voxel pattern similarity between the fixation modulation effect and subsequent memory effect in HPC and PPA ROIs are greater in the scene *free-viewing* than the *fixed-viewing* condition. Ventral brain surface views of voxel-wise fixation modulation effects (left) and subsequent memory effect (right) showing the ventral and posterior medical temporal lobe regions are each associated with both the number of gaze fixations and subsequent memory performance. Voxel-wise fixation modulation and subsequent memory effect in HPC and PPA at the group level are embedded to illustrate the similarity between the two effects in the *free-viewing* condition. **D**. The number of fixations predicted activation in PPA bilaterally after controlling for subsequent memory effect. **E**. Activation of a cluster in the PPA and HPC bilaterally was positively predicted by the number of gaze fixations after controlling for subsequent memory effect (*p*=.005, with 10-voxel extension, no corrections). **F**. & **G**. Brain surface views for the prediction of the number of gaze fixations (**F**) and subsequent memory (**G**) for the scenes under *free viewing* after controlling for shared variability between the two predictors. **H**. The whole brain cross-voxel pattern similarity between the fixation modulation and subsequent memory is higher in the *free-* than *fixed-viewing* conditions for scenes, with or without controlling for the shared variability between the two variables. A horizontal section of the whole brain voxel-wise fixation modulation and subsequent memory effect at the group level are embedded to illustrate the similarity between the two effects in the *free-viewing* condition. Note: Surface and section view **B, E, F**, and **G** are presented at *p*<.005, 10-voxel extension with no corrections. For the brain surface views, L indicates left hemisphere and R the right hemisphere. For **A, C, D**, and **H**, + = *p*<.10; *=*p*<.05; **=*p*<.005; ***= *p*<.001, one-tailed t test.

To confirm that the PPA and HPC were modulated by both trial-wise number of fixations and subsequent memory, we did a conjunction analysis in which we thresholded the gaze fixation and subsequent memory modulation effects at *p* = 0.05, with 50-voxel extension (no corrections) and examined the overlap. This revealed a conjunction effect with *p* < 0.0025, 50 voxel-extension (no correction) in which both the PPA and a cluster of voxels in the HPC showed stronger activation when scenes were viewed with more fixations *and* were subsequently remembered (Figure 6B).

Although we found that the two behavioral variables (the number of fixations and subsequent memory) had shared neural variance, the conjunction analysis cannot test the extent to which there is a similar pattern of brain activity associated with the two behavioral variables. To confirm that the trial-wise number of fixations and subsequent memory were also related at the brain level, we conducted a pattern similarity analysis in which the cross-voxel (within the HPC and PPA) subsequent memory effect pattern of neural activity was correlated with the cross-voxel gaze fixation modulation effect pattern of neural activity. If visual exploration (as indexed by the number of fixations) is indeed associated with memory formation at the brain level in the *free-viewing* condition for scenes, the cross-voxel pattern of the trial-wise number of fixations modulation effects should be correlated with the cross-voxel brain activity pattern that reflects subsequent memory performance. As shown in Figure 6C, consistent with our hypothesis, the pattern similarity between the gaze fixation and subsequent memory modulation effects was significantly larger than zero for the left and right HPC (*t* = 3.30 and 3.92, *p* < .005 and .001, respectively) and the left and right PPA (*t* = 5.46 and 4.55, *p* < .001) in the scene *free-viewing* condition. The similarity was not significant in the scene *fixed-viewing* condition (*t* = .13 ∼ 1.63, *p* = .90 ∼ .11 for the left and right HPC and PPA). Moreover, the similarity between the subsequent memory and fixation modulation effects was significantly stronger in the *free-* versus *fixed-viewing* condition (*t* = 2.54 and 2.36, *p* = .008 and .012 for the left and right HPC; *t* = 3.08 and 1.95, *p* = .002 and .03 for the left and right PPA, one-tailed; Figure 6C). Thus, in the *free-viewing* condition, the trial-wise number of fixations and subsequent memory performance shared variability at both the behavioral and the brain (individual voxel) level. When visual exploration was constrained in the *fixed-viewing* condition, the association between gaze fixations and memory at both levels was disrupted.

We further examined whether the number of fixations predicted brain activity in the HPC and PPA even after controlling for effects of subsequent memory. We simultaneously entered the two modulator regressors (i.e., trial-wise number of fixations and subsequent memory) to the same parametric modulation analysis and examined the modulation effects of each regressor without the contribution of the other. Our parametric modulation ROI analysis showed that a greater number of gaze fixations indeed predicted stronger activation in the left and right PPA (*t* = 3.96 and 5.35, *p* <.001. Figure 6D). The effect was not significant for the left and right HPC (*t* = .29 and .73, *p* =.77 and .47, respectively) when the whole HPC ROIs were used. However, as shown in Figure 4B, the modulation effect of the number of fixations did not spread to the whole HPC. We therefore did a similar small volume correction analysis using the HPC template masks to correct for multiple comparisons, and thresholded the whole brain voxel-wise results with *p* = .005, 10-voxel extension (without corrections). The result revealed a cluster of voxels in both the left and right HPC that survived family-wise error correction at *p*_fwe-corr_ = .05 (for the left HPC: *p*_fwe-corr_ = .023, *p* < .0001, voxel size = 105 voxels with the peak voxel location = [−20, −32, −2]; for the right HPC: *p*_fwe-corr_ = .012, *p* < .0001, voxel size = 140 voxels with the peak voxel location = [28, −34, 4]). Brain section and surface images in Figure 6E and 6F illustrate the gaze fixation modulation effects in the PPA and HPC, as well as other brain regions, that occur above and beyond (and cannot be merely attributed to) subsequent memory effects.

For completeness, we examined the subsequent memory effects after controlling for the modulation effect of the number of gaze fixations. Numerous brain regions, including the ventral and posterior medial temporal lobe regions, were modulated similarly by visual exploration and subsequent memory performance for the scenes under *free-viewing* conditions, even after the shared variance between the two variables was excluded (Figure 6G). To quantify the similarity between the two modulation effects at the whole-brain level, we calculated the modulation pattern similarity measure across all voxels in the brain between the fixation and subsequent memory effect using the same method as in Figure 6C. The similarity measure in the scene *free-viewing* condition was significantly larger than zero (*t* = 2.93, *p* =.006, one-tailed; Figure 6H), and significantly greater than the similarity measure in the *fixed-viewing* condition (*t* = 2.33, *p* =.013, one-tailed). Similarly, we calculated the whole-brain cross-voxel similarity between the two modulation effects when the shared variance between the two behavioral measures was retained. The similarity measure was again stronger in the *free-* than *fixed-viewing* condition (*t* = 3.70, *p* <.001, one-tailed). The similarity measures were also greater when the shared variance was retained than when excluded (*p* < .001; Figure 6H), confirming that the shared variance at the behavioral level between the number of gaze fixations and subsequent memory was also reflected at the whole-brain cross-voxel level.

## Discussion

There is an intimate link between oculomotor behavior and memory^19^. The current study shows that robust relationships exist between naturalistic visual exploration and memory at both the behavioral and brain levels. Such findings lend strong support to the notion that gaze fixations provide the requisite units of information on which hippocampal binding processes operate to create new memories. This work provides the first non-invasive, comprehensive, and direct evidence of an oculomotor-memory system dependency in humans.

Visual exploration is positively related to recognition memory^8, 9^, and the restriction of viewing leads to a reduction in subsequent memory performance^10, 11^. Findings from neurophysiology and neuroimaging have shown that oculomotor behavior modulates neural responses in regions that are critical for memory^20, 21^. When considered together, we predicted causal relationships between visual exploration, neural activity, and memory, such that restricting visual exploration would decrease neural responses in the HPC, broader medial temporal lobe, and along the ventral visual stream; change the functional connectivity between the HPC, PPA, and other cortical regions; and reduce subsequent memory. Here, we replicated our prior work using face stimuli^12, 13^ by showing that neural activity in the HPC, as well as in the PPA, were positively associated with the number of gaze fixations during free viewing of scenes. Critically, experimentally reducing visual exploration led to a reduction in neural activity all along the ventral visual stream, and up into the PPA and HPC, and reduced subsequent recognition memory. Gaze fixation modulation effects were not significant in the HPC or PPA for scenes under *fixed-viewing*, or for either of the scrambled picture conditions for which little information is available for extraction.

Given its well-established role in the processing of scenes^17, 22^, we interrogated the functional connectivity of the PPA. When compared to *fixed-viewing, free-viewing* was associated with increased functional connectivity with the HPC, as well as with lateral and medial occipital cortex, and increased negative functional connectivity with regions such as the inferior parietal lobule, precuneus, anterior cingulate, lateral PFC, angular gyrus, and supramarginal gyrus. Reducing visual exploration therefore led to decreases in the relative strength of the functional connections across the network, and may have reduced information exchange between the oculomotor and medial temporal lobe networks^4, 15^, resulting in declines in subsequent recognition memory. Thus, naturalistic viewing may coordinate responses (through correlated or potentially anti-correlated activity) across a broad network that includes regions responsible for the cognitive control of eye movements, perceptual processing of visual information, updating of spatial information, and memory^15,23,24^.

The neural pattern of gaze modulation effects was similar to that of subsequent memory effects. The conjunction analysis confirmed that there were clusters in the PPA and the HPC where activity was predicted by the number of gaze fixations as well as subsequent memory. The cross-voxel brain activation patterns in the HPC, PPA, and even the whole brain that were associated with the number of gaze fixations were more similar to those associated with subsequent memory when scenes were encoded under *free-viewing*, compared to when scenes were encoded under *fixed-viewing* conditions. Importantly, clusters of neural activity in the HPC and PPA that were modulated by gaze fixations under *free-viewing* were evident even after controlling for the effect of subsequent memory. Together, these findings suggest that gaze fixations are related to the perceptual processing and binding functions of ventral visual stream, and HPC, respectively, above and beyond whether the resultant memory representations are available for conscious introspection. This is consistent with the purported role of the HPC in relational binding, in which elements are bound across space and time into a lasting, flexible representation, and may guide ongoing behavior in the moment, even in the absence of conscious awareness for the information contained therein^25–29^. The added effect of gaze fixations on neural activity, beyond the effects of subsequent explicit recognition, is also consistent with work in which viewers, in the absence of visual input, follow distinct scanpaths of other participants that had been elicited by viewing either faces or houses, show differential activation the fusiform face area and PPA, respectively^30^. It is likely that lifelong experience viewing faces and houses creates stored exemplars of scanpaths that represent stimulus-specific viewing. Although future work is needed to explore how neural activity relates to the reproduction of specific scanpaths, the present findings suggest that, at the very least, gaze fixations contribute information that supports the development of memory. The sequence and location of those gaze fixations comprise the scanpath that may be part-and-parcel of any resultant memory trace ^31^, even in the absence of conscious recognition for the viewers’ own scanpath^32^ or for the previously studied stimulus.

Restricting visual exploration had a general effect in reducing associated neural activity, as neural responses were greater along the ventral visual stream and into the medial temporal lobe under *free-viewing* conditions for scenes and for scrambled images alike. This may suggest that eye movements simply have a faciliatory effect on neural activity, such that gaze fixations create a feedforward sweep of neural responses that prime the brain to efficiently process upcoming visual input. However, differences in neural activity for the *free-* versus *fixed-viewing* contrast were larger for scenes than for the scrambled images. Moreover, the cross-voxel modulation pattern of subsequent memory resembled that of visual exploration, which was stronger when scenes were viewed under the *free-* versus *fixed-viewing* condition. These findings suggest that visual exploration is related to the processing and encoding of informational content^5, 6^. An alternative account is that the reduction in neural responses during *fixed-* versus *free-viewing* reflects a decrease in saccadic suppression that would otherwise occur with movements of the eye^33^. However, research points to an associated *reduction* of neural responses in visual processing regions such as V4 that occurs with eye movements^34^. Here, the opposite occurred: there was an increase in neural activity along the visual processing stream associated with *free-viewing*, the condition for which greater saccade suppression should occur due to increased numbers of gaze fixations relative to *fixed-viewing*.

Another consideration for interpreting the present findings is whether *fixed-viewing* resulted in an increase in working memory demands in an effort to remain continually fixated that, in turn, altered the level of neural engagement and negatively affected memory. Prior work has shown that maintaining central fixation does not seem to increase working memory demands, as there was no reduction in n-back performance under *fixed-* versus *free-viewing* conditions^35^. By contrast, performing a finger tapping task did interfere with n-back performance relative to *free-viewing*, as evidenced by a decrease in accuracy and an increase in response times^35^. Thus, restricting visual exploration here likely resulted in a decrease in memory performance due to an inability to bind extracted visual information into a lasting memory representation, rather than due to an increased demand on working memory capacity.

We urge methodological caution for neuroimaging (fMRI, electroencephalography, magnetoencephalography) studies that require central fixation in an effort to reduce motion artifacts in the data; the findings here demonstrate how restriction of viewing behavior changes the amount of engagement within neural regions and across functional networks, and changes subsequent cognitive performance. This is not to say that central fixation would influence every study in such a manner. Viewing manipulations may have little to no effect when tasks employ simple stimuli and/or use simple task demands. However, maintaining central fixation may adversely affect the pattern of data that is observed when more complex stimuli are used, as seen here, or when the participant engages in more complex tasks (for further discussion, see ^19^).

The present findings also call for future research that examines the level or type of information that is carried via gaze fixations on which the HPC (and broader medial temporal lobe) operates. Multiple theoretical accounts of hippocampal function purport that the HPC receives already-parsed information regarding objects, spatial locations, and temporal ordering for the purpose of then binding those elements into a coherent, lasting memory representation^26,28,36^. However, oculomotor research typically considers the allocation of gaze fixations for the purpose of extracting information regarding lower-level features, such as luminance and contrast^7^. Evidence, especially the findings reported here, suggests that the HPC may bind visual information online, across gaze fixations^12,37,38^. How, then, do we bridge this theoretical divide and understand the unit of information that gaze fixations carry to which the HPC is sensitive? Through its movements over space and time, gaze fixations inherently provide information regarding the spatial and temporal arrangements of distinct elements, and a plethora of research shows that cells in the HPC respond preferentially to particular places or temporal orders^39–41^. Alternatively, gaze fixations may serve to organize the sequential activity of neuronal assemblies in the HPC, from which properties of space and time may be derived^42^. While work remains to specifically address these issues, the present research provides key causal evidence supporting the notion that the oculomotor system is a unique effector system for memory^1^ such that gaze fixations provide the fundamental units of information on which the binding functions of the hippocampus operate.

## Online Methods

### Participants

Thirty-six healthy young adults (22 females; age: *M* = 23.58 years, *SD* = 4.17; education: *M* = 16.27 years, *SD* = 1.8) participated in exchange for monetary compensation. All participants had normal or corrected-to-normal vision (including color vision), and none had any neurological or psychological conditions. Participants were recruited from the University of Toronto and surrounding Toronto area community. All participants provided written informed consent. The study was approved by the Research Ethic Board at Rotman Research Institute at Baycrest Health Sciences.

### Stimuli

Stimuli included twenty-four color scene images (500 × 500 pixels; viewing angle: 7.95 × 7.95 degree) for each of 36 semantic scene categories (e.g., living room, arena, warehouse, etc.), for a total of 864 images, were used in the present study. Half of the scene images were from a previous study by Park et al.^43^. The other half of the scene images were collected from Google Image using the 36 categories as the search terms. These newly collected images matched the features of the original images by Park et al^43^. The 36 categories of scenes varied along two feature dimensions: the size of scene space and the clutter within the scene. Each feature dimension (size, clutter) had 6 levels in a balanced factorial design such that each level of one feature contained 6 levels of the other feature, resulting in 36 unique feature level combinations (for details, see Park et al^43^).

The 24 scene images in each scene category were randomly divided into 3 groups with 8 images in each group. One group of images (8 images/category x 36 categories = 288 images) was used for the *free-viewing* encoding condition, one group (288 images) for the *fixed-viewing* encoding condition, and one group (288 images) served as the lure images during the retrieval task. The assignment of the 3 groups of images to these experimental/stimulus conditions was counterbalanced across participants. Low-level features such as luminance and contrast were equalized across the three groups of images using a Matlab script based on the SHINE toolbox. Color histogram and spatial frequencies were calculated using Natural Image Statistical Toolbox for MATLAB^44^ and also balanced across the 3 groups of images.

The 8 images of each scene category in the free- and fixed-viewing conditions were randomly assigned to 8 fMRI encoding runs). Each run had 36 images (one scene from each category) that were viewed under *free-viewing* instructions and 36 images (one scene from each category) that were viewed under *fixed-viewing* instructions. Then, images from two randomly selected runs were scrambled, using 6 levels of tile sizes (i.e., 5, 10, 25, 50, 100, and 125 pixels) to resemble the cluster feature of the scene pictures, to create two runs of scrambled images. Similarly, two images from each scene category that were to be used as lures in the retrieval phase were also scrambled to provide scrambled lure images for the retrieval task.

To summarize, 6 images x 36 categories = 216 scene images and 2 images x 36 categories = 72 scrambled images were used in the *free-viewing* encoding condition; the same number of images was used in the *fixed-viewing* encoding condition. All of the images from the encoding phase, together with an additional 6 images x 36 categories = 216 scene images and 2 images x 36 categories = 72 scrambled images, were used in the retrieval task.

The feature levels of the scene images (i.e., the size of scene space and the clutter within the scene) were fully balanced across the groups of images used during *free-viewing* and *fixed-viewing* encoding and retrieval images. For our current purposes, the focus was on the neural differences between the free viewing and fixed viewing encoding conditions; the feature level manipulation (size, clutter) was not considered in the current analyses.

### Procedure

As mentioned above, the encoding task had 8 runs; 6 runs contained images of scenes, and 2 runs contained scrambled images. Each run had 72 images; 36 of which were studied under free-viewing instructions, and 36 of which were studied under fixed-viewing instructions, with one image from each of the 36 scene categories for each viewing condition. The order of the 6 scene and 2 scrambled image runs was randomized for each participant.

During each encoding trial (see Figure 1), a fixation cross “+” was first presented for 1.72 – 4.16 seconds (exponential distribution with mean = 2.63 seconds), and participants were asked to fixate their eye gaze on the cross. The location of the cross was randomly determined within a radius of 100 pixels around the center of the scene image. The cross “+” could be presented in either a green or red color. After the fixation cross, a scene image was presented for 4 seconds. When the cross was green, participants were to freely explore the scene or scrambled image that followed in order to encode as much information as possible (i.e., the *free-viewing* condition). When the cross was red, participants were required to keep their eye gaze at the location of the fixation cross throughout the presentation of the image that followed (i.e., the *fixed-viewing* condition). The task for each run was 500 seconds long, with 10 seconds and 12.4 seconds added to the beginning and end of the task, respectively. The trials from the two viewing conditions were pseudo-randomized to obtain an adequate design efficiency by choosing the design with the best efficiency from 1000 randomizations using Matlab code by Spunt^45^.

### Structural and functional MRI

A 3T Siemens MRI scanner with a standard 32-channel head coil was used to acquire structural and functional MRI images. T1-weighted high-resolution MRI images for structural scans were obtained using a standard 3D MPRAGE (magnetization-prepared rapid acquisition gradient echo) pulse sequence (170 slices; FOV = 256 × 256 mm; 192×256 matrix; 1 mm isotropic resolution, TE/TR = 2.22/200ms, flip angle = 9 degrees, and scan time = 280 s). For the fMRI scan, the BOLD signal was assessed using T2*-weighted EPI acquisition protocol with TR = 2000 ms, TE = 27 ms, flip angle = 70 degrees, and FOV = 191 × 192 ms with 64 × 64 matrix (3 mm x 3 mm in-place resolution; slice thickness = 3.5 mm with no gap). Two hundred and fifty volumes were acquired for each fMRI run. Both structural and functional images were acquired in an oblique orientation 30° clockwise to the anterior–posterior commissure axis. Stimuli were presented with Experiment Builder (Eyelink 1000; SR Research) back projected to a screen (projector resolution: 1024×768) and viewed with a mirror mounted on the head coil.

### Eyetracking

Participants’ eye movements were recorded using an MRI-compatible eye tracker (Eyelink 1000; SR Research) with a sampling rate of 1000 Hz and spatial resolution of 1°. Calibration was done using the built-in EyeLink 9-point calibration procedure at the beginning of the experiment. Drift correction was performed between trials when necessary to ensure good tracking accuracy. Fixations and saccades were categorized using Eyelink’s default eye movement event parser. Specifically, a velocity threshold of 30°/s and acceleration threshold of 8000°/s were used to classify saccades (saccade onset threshold = 0.15°). Events not defined as saccades or blinks were classified as fixations. The number of fixations that participants made when they encoded each scene was calculated and exported using the EyeLink software Data Viewer.

### fMRI data preprocessing

SPM 12 (Statistical Parametric Mapping, Welcome Trust Center for Neuroimaging, University College London, UK; https://www.fil.ion.ucl.ac.uk/spm/software/spm12/ Version: 7487) in the Matlab environment (The MathWorks, Natick, USA) was used to process the functional images. Following the standard SPM 12 preprocessing procedure, slice timing was first corrected using sinc interpolation with the midpoint slice as the reference slice. Then, all functional images were aligned using a six-parameter linear transformation. Next, for each participant, functional image movement parameters obtained from the alignment procedure, as well as the global signal intensity of these images, were checked manually using the freely available toolbox ART (http://www.nitrc.org/projects/artifact_detect/) to detect volumes with excessive movement and abrupt signal changes. Volumes indicated as outliers by ART default criteria were excluded later from statistical analyses. Anatomical images were co-registered to the aligned functional images and segmented into white matter (WM), gray matter (GM), cerebrospinal fluid (CSF), skull/bones, and soft tissues using SPM 12 default 6-tissue probability maps. These segmented images were then used to calculate the transformation parameters mapping from the individuals’ native space to the MNI template space. The resulting transformation parameters were used to transform all functional and structural images to the MNI template. For each participant, the quality of co-registration and normalization was checked manually and confirmed by two research assistants. The functional images were finally resampled at 2×2×2 mm resolution and smoothed using a Gaussian kernel with an FWHM of 6 mm. The first five fMRI volumes from each run were discarded to allow the magnetization to stabilize to a steady state, resulting in 245 volumes in each run.

### fMRI analysis

#### Activation differences: Free-vs. fixed-viewing

We used SPM 12 to conduct the first (i.e., individual) level whole brain voxel-wise General Linear Model (GLM) analysis to examine brain activation differences between the *free-viewing* and *fixed-viewing* encoding conditions for both the scenes and scrambled pictures. In this event-related design, we separately convolved the onset of trials in the free-viewing and fixed-viewing condition with the canonical hemodynamic function (HRF) in SPM 12, which served to be the two main regressors of interest. Because there were runs of scenes, and runs of scrambled images, we had 4 regressors of interest: free-viewing of scenes, fixed-viewing of scenes, free-viewing of scrambled pictures, and fixed-viewing of scrambled pictures. motion parameters (6 from SPM realignment, 1 from ART processing), as well as outlier volumes, were added as regressors of no interest. Default high-pass filter with cut-off of 128 s was applied. A first-order autoregressive model AR(1) was used to account for the serial correlation in fMRI time-series in the restricted maximum-likelihood estimation of the GLM. To examine the main question of whether brain activation was different for the *free-* and *fixed-viewing* condition, we constructed a *t* contrast to directly compare the two conditions ([*free-viewing – fixed-viewing*]) for each run and then averaged all 6 scene runs (note that one participant only had 5 scene runs). We did a similar analysis to compare the *free-* and *fixed-viewing* conditions for the scrambled pictures. We also constructed a *t* contrast to compare the viewing effect (i.e., the difference between the two viewing conditions) between the two types of stimuli (scenes versus scrambled pictures).

Because we had specific *a priori* brain regions of interest, i.e., the hippocampus (HPC) and parahippocampal place area (PPA), at the second (i.e., group) level we used a region of interest (ROI; For ROI definition, see ROI section) analysis approach. Specifically, we extracted the mean beta estimates within each ROI for each contrast (effect) and tested the ROI effect using one-sample *t-*tests. Because the HPC has an elongated shape and it is likely that not all segments along the HPC longitudinal axis would show the same effect, we also examined the voxel-wised results within HPC, using all voxels in the HPC masks to perform the family-wise multiple comparison error correction (Threshold: *p*_fwe-corr_ = .05). SPM 12’s small-volume-correction procedure and the HPC MIN template masks were used in this analysis.

#### Parametric modulation analysis

In order to replicate our prior work^12, 13^ with face stimuli, we conducted a parametric modulation analysis in SPM 12 for the *free-viewing* scene condition to examine whether increases in the number of gaze fixations was associated with stronger activation in the brain’s memory and perceptual processing regions. Specifically, in addition to regressors in the design matrix described in the previous section, we added a linear parametric modulation regressor that consisted of the trial-wise number of fixations convolved with the canonical HRF. Although we added this regressor for all 4 conditions in the design matrix, the focus here is on the *free-viewing* scene condition based on our prior findings^12, 13^. The effect of the fixation modulation regressor in the *free-viewing* condition was averaged across all scene runs and carried to the second (i.e., group) level ROI analysis. The same approach for the second level analysis, as mentioned earlier, was used for this analysis.

To understand the extent to which there was similar modulation of brain activity by the number of fixations and by subsequent memory, we also used parametric modulation analysis to examine subsequent memory effects. We coded subsequent recognition memory for each encoding trial based on participants’ hit/miss response and confidence. Specifically, we assigned 2 points to stimuli that were correctly recognized as previously viewed with high confidence, 1 point for those correctly recognized with low confidence, 0 points for previously viewed images incorrectly endorsed as “new” with low confidence, and −1 point for previously viewed images that were incorrectly endorsed as “new” with high confidence to construct the linear parametric modulator. The design matrix, contrasts, and the first and second level analysis procedures were identical to those used in the fixation modulation analysis, save for the use of different modulation regressors.

To test whether the modulation effect of the trial-wise number of fixations on HPC and PPA activity in the *free-viewing* scene condition was associated with the brain activity that supports subsequent memory, we did a conjunction analysis in which we thresholded the voxel-wise fixation modulation effects and subsequent memory effects at *p* = 0.05, with 50-voxel extension (no corrections) and examined the conjunction (i.e., overlap) map between the two modulation effects. This resulted a conjunction effect map with *p* < 0.0025, 50 voxel-extension (no correction).

In addition to focusing on the overlap between two modulation effects at the mean level, we also investigated cross-voxel similarity between the subsequent memory effect and the modulation effect of the trial-wise number of fixations in the HPC and PPA. This was done to test whether the brain activity associated with two behavioral variables (i.e., trial-wise number of fixation and subsequent memory) also share similar brain activity pattern. Specifically, for each participant, we extracted first-level subsequent memory and fixation modulation effect (i.e., the first level GLM beta estimates) from each voxel in the HPC and PPA ROIs. These voxel-wise beta values from the two parametrical modulation analyses were vectorized for each ROI. Then, Pearson correlation was calculated between the two vectors and then Fisher’s Z transformed to reflect the cross-voxel pattern similarity between the subsequent memory effect and the modulation effect of the number of fixations. We obtained the similarity measure for both the scene *free-viewing* and *fixed-viewing* conditions and compared them at the group level. If the brain activity pattern modulated by the number of fixations are indeed important for supporting the subsequent memory, we should observe greater cross-voxel similarity in the *free-viewing* than the *fixed-viewing* condition. This would provide further evidence supporting that visual exploration and memory formation share similar brain mechanisms.

Finally, we examined whether the trial-wise number of fixations still predicted activity in the HPC after controlling for the effect of subsequent memory. We reasoned that if visual exploration is generally important for hippocampal processes, irrespective of subsequent conscious awareness, the trial-wise number of fixations should still predict the brain activity in the medial temporal lobe after the shared variance with subsequent memory was partialed out. To test this hypothesis, we simultaneously entered the two parametric modulation regressors, i.e., trial-wise subsequent memory and number of fixations, to the same parametric modulation analysis. Then, we examined the modulation effects of the number of fixations without the contribution of subsequent memory. In this analysis, we used the identical first and second level analysis procedure as mentioned above (except for using two parametric modulators) and focused on the *free-viewing* condition for scenes.

#### PsychoPhysiological Interaction (PPI) analysis

To investigate how brain connectivity may differ between the *free*- and *fixed-viewing* condition, we conducted a generalized psychophysiological interaction analysis (gPPI) ^46, 47^ using the PPAs as seed regions. We chose the PPAs as the seed regions for two reasons. First, the PPA is a key brain region that supports the perceptual processing of scenes, and it can be well localized in this task. Because the HPC is an elongated structure and different regions of the HPC along its longitudinal and sectional axis may be differentially involved in this task, it is not ideal to use the whole HPC as the seed region. Second, the PPA is closely connected with visual processing regions earlier in the ventral visual stream, as well as the HPC, which is critical for memory^48, 49^. Using the PPA as the seed region allowed us to examine how the connectivity among perceptual processing and memory regions in the brain can be modulated by different viewing conditions. We conducted the gPPI analysis using CONN toolbox v.18 (https://www.nitrc.org/projects/conn) in Matlab^50^. First, all preprocessed functional images using SPM 12 were further preprocessed to eliminate signals that may affect the connectivity analysis. BOLD signal from the WM and CSF were used to regress out non-specific variance from the fMRI time series using a principal component-based noise correction method ^51^ as implemented in CONN toolbox. The first five principal components extracted from the WM and CSF were used in this noise removal procedure. Motion parameters obtained from the motion correction procedure were also used to regress out potential head movement effects. Slow fMIR signals were filtered out using .008 Hz high-pass filter. The data were also de-spiked using a hypobolic tangent function to reduce the impact of any potential outliers in the fMRI time series. After these further clean-up procedures, fMRI time series, averaged from the left and right PPA, were extracted from each encoding run. An interaction term was formed between the PPA time series and all other task condition regressors (which were convolved with the canonical HRF). Then, the original task condition regressors, PPA time series, and the interaction regressor were entered in the same GLM analysis. The contrast of the effect of the interaction regressor between the scene *free-viewing* and *fixed-viewing* conditions, reflecting the differences in PPA connectivity with other brain regions due to differences in viewing instructions, was the focus of this analysis. The results from all scene runs were averaged and carried to the second level analysis. The same approach for the second level analysis, as mentioned earlier, was used for this analysis.

### ROI definition

HPC ROIs were defined anatomically: The bilateral HPC masks in individual participants’ native space were first obtained using FreeSurfer *recon-all* function, version 6.0 (http://surfer.nmr.mgh.harvard.edu.myaccess.library.utoronto.ca) ^52^. Then, the same normalization parameters obtained from the SPM normalization procedure were used to transform these hippocampal masks into MNI normalized space.

PPA ROIs were defined functionally: we first contrasted the scene and scrambled pictures conditions, collapsed over the *free-* and *fixed-viewing* conditions, at the individual-level analyses. Then, at the group level, we localized the maximumly activated cluster in the bilateral parahippocampal cortex at the threshold of *p*_fwe-corr_ = .05 (family-wise error multiple comparison correction). The MNI coordinates for the peak activation of the contrast were [32, −34, −18] for the right PPA and [−24, −46, −12] for the left PPA. The left and right PPA mask contained 293 and 454 voxels respectively (see Supplementary Figure S1).

### Statistical thresholding

The threshold for statistical significance was set at *p* = .05 for the ROI analyses when the mean value of the estimated effect was obtained. For the whole HPC, when the results were not significant, *p*_fwe-corr_ = .05 (family-wise error multiple comparison correction) was used within the HPC (i.e., small volume correction) to examine whether voxel clusters in the HPC showed significant effects. We also present the whole-brain voxel-wise results thresholded at *p* = .005 with 10 voxel extension (uncorrected) to facilitate future meta-analysis ^53^. The automated anatomical labeling (AAL) toolbox ^54^ was used to identify the anatomical labels for regions that showed significant effects, which are reported in Supplementary Tables.

### Post-fMRI retrieval task

After the fMRI encoding task, participants were given a 60-minute break before they began a retrieval task in a separate testing room. All scenes and scrambled images from the fMRI encoding task (36 images/condition/run x 2 conditions x 8 runs = 576 images) were tested in the retrieval task. In addition, 288 images that were not used in the encoding task (6 images x 36 categories =216 scene images and 2 images x 36 categories = 72 scrambled images) were also included as lures. The 864 images were divided into 6 blocks.

For each retrieval trial, a fixation cross “+” was presented for 1.5 seconds. For the previously viewed images (i.e., images from the encoding phase), the cross “+” was presented at the same location as it was during the encoding task. Following the presentation of the fixation cross, the retrieval image was presented on the screen for 4 seconds. Participants were given 3 seconds to indicate whether the image was a “new” or “old” (i.e., previously viewed during the fMRI task) image using 4 keys on the keyboard: z – high confidence “old”, x – low confidence “old”, n – high confidence “new”, and m – low confidence “new.” The response instruction was shown on the screen during the time window in which participants made their response. During the retrieval task, participants’ eye movements were recorded using the Eye-link II head mounted infrared camera system (SR Research Ltd., Oakville, ON, Canada) at a sampling rate of 500 Hz. However, the eye movement data from the retrieval task were not relevant for our current focus and are not discussed further here.

To measure recognition memory, we took participants’ response confidence into consideration and assigned 2 points to stimuli that were correctly recognized as previously viewed with high confidence, 1 point for those correctly recognized with low confidence, 0 points for previously viewed images incorrectly endorsed as “new” with low confidence, and −1 point for previously viewed images that were incorrectly endorsed as “new” with high confidence. This recognition memory measure was then used to investigate the behavioral consequence of the manipulation of viewing condition, the relationship between the trial-wise number of fixations and memory performance, and brain regions supporting memory through subsequent memory analyses.

## Acknowledgments

We would like to thank Arber Kacollja, Sarah Berger, Ling Li, Mandy Ding, Veena Sanmugananthan, Junghyun Nam, Mariam Aziz, and Jordana Wynn for their help at different stages of this research project. This research was supported by a Vision: Science to Applications (VISTA) postdoctoral fellowship awarded to ZXL, funding from the Natural Sciences and Engineering Research Council of Canada awarded to JDR and to RSR, and the Canadian Institutes of Health Research awarded to JDR.

