## Supplementary Material for "Restricting visual exploration directly impedes neural activity, functional connectivity, and memory"

**Table S1.** Mean memory performance and standard deviation (*SD*) in the *free-viewing* and *fixed-viewing* conditions.

| | Scenes | | | | Scrambled | | | |
|---|---|---|---|---|---|---|---|---|
| | recognition memory | corrected accuracy | hit rate | false alarm rate | recognition memory | corrected accuracy | hit rate | false alarm rate |
| Free viewing | .46 | 11.47% | 46.71% | 35.18% | .22 | 0.74% | 34.85% | 34.10% |
| | (.32) | (8.25%) | (14.99%) | (14.00%) | (.30) | (7.41%) | (19.91%) | (20.10%) |
| Fixed viewing | .31 | 5.06% | 40.43% | 35.18% | .22 | 0.75% | 34.55% | 34.10% |
| | (.33) | (5.03%) | (15.25%) | (14.00%) | (.32) | (8.09%) | (19.35%) | (20.10%) |

***Notes*:**

a. Recognition memory was calculated by assigning 2 points to stimuli that were correctly recognized with high confidence, 1 point for those correctly recognized with low confidence, 0 points for previously viewed images endorsed as "new" with low confidence, and -1 point for previously viewed images endorse as "new" with high confidence. A 2 x 2 repeated ANOVA (free-/fixed-viewing by scenes/scrambled pictures) showed that recognition memory was better for scenes than for scrambled pictures ($F(1,35) = 6.92$, $p < .013$), and higher for the images studied under *free-viewing* versus fixed-viewing instructions ($F(1,35) = 27.23$, $p < .0001$). This latter difference was significant only for the scenes (Interaction effect: $F(1,35) = 27.85$, $p < .0001$);

b. Corrected accuracy = hit rate – false alarm rate; The false alarm rate is the same for the free and fixed viewing condition because the viewing condition is not relevant for the new pictures in the retrieval task. Similarly, a 2 x 2 repeated ANOVA (free-/fixed-viewing by scenes/scrambled pictures) showed that corrected accuracy was higher for scenes than for scrambled pictures ($F(1,35) = 27.91$, $p < .0001$), and higher for the images studied under *free-viewing* versus fixed-viewing instructions ($F(1,35) = 21.35$, $p < .0001$). This latter difference was significant only for the scenes (Interaction effect: $F(1,35) = 22.91$, $p < .0001$);

c. We also examined whether the trial-wise number of fixations predicted recognition memory. Specifically, we ran a linear regression analysis for each condition in each run using the trial-wise number of fixations to predict recognition memory. The regression coefficient was averaged across runs for each participant. These coefficients from all participants were then subjected to independent t-tests to determine whether the mean was significantly higher than zero. This approach is identical to the two-level fMRI analysis approach used in this study.

d. We note that memory accuracy may have been relatively low in this experiment. This is likely due to the following factors: 1) a 1-hour delay between the encoding and retrieval phases, 2) old and new scenes were selected from the same sematic categories, and 3) large numbers of stimuli were used.

**Table S2.** Brain regions that showed stronger or weaker activation during *free-viewing*, compared to *fixed-viewing*, of scenes.

| Anatomical areas | Cluster size | *t* value | *p* value | MNI coordinates | | |
|---|---|---|---|---|---|---|
| | | | | x | y | z |
| **Free-viewing > fixed-viewing** | | | | | | |
| Lingual_L_R* | 45470 | 12.15189 | 2.07E-14 | 22 | -44 | -8 |
| Frontal_Inf_Orb_R | 295 | 6.565433 | 7E-08 | 32 | 32 | -16 |
| Frontal_Inf_Tri_R | 352 | 4.642333 | 2.35E-05 | 56 | 24 | 28 |
| Insula_L | 186 | 4.416027 | 4.61E-05 | -32 | -22 | 20 |
| Cingulum_Ant_R | 76 | 4.322811 | 6.08E-05 | 6 | 2 | 26 |
| Postcentral_L | 80 | 3.582199 | 0.000513 | -62 | -10 | 34 |
| Postcentral_R | 48 | 3.560667 | 0.000544 | 62 | -6 | 36 |
| Frontal_Sup_L | 24 | 3.549678 | 0.000561 | -20 | 36 | 52 |
| Cerebelum_Crus2_R | 30 | 3.517169 | 0.000614 | 8 | -78 | -36 |
| Postcentral_R | 46 | 3.298407 | 0.001119 | 48 | -6 | 30 |
| Insula_R | 19 | 3.262465 | 0.001233 | 34 | -20 | 16 |
| Cingulum_Mid_R | 19 | 3.172249 | 0.00157 | 8 | -10 | 42 |
| Frontal_Sup_L | 14 | 3.165734 | 0.001598 | -16 | 64 | 18 |
| Temporal_Pole_Sup_L | 14 | 3.145887 | 0.001685 | -46 | 16 | -16 |
| **Fixed-viewing > free-viewing** | | | | | | |
| Parietal_Inf_R | 1685 | 9.128221 | 4.36E-11 | 56 | -52 | 46 |
| Parietal_Inf_L | 729 | 7.037532 | 1.71E-08 | -54 | -56 | 42 |
| Frontal_Inf_Oper_R | 373 | 6.334046 | 1.41E-07 | 50 | 4 | 18 |
| Cingulum_Post_R | 584 | 5.485443 | 1.84E-06 | 14 | -36 | 18 |
| Pallidum_R | 34 | 4.874105 | 1.17E-05 | 16 | 6 | -4 |
| Frontal_Mid_R | 478 | 4.441552 | 4.28E-05 | 40 | 28 | 38 |
| Insula_R | 61 | 4.395148 | 4.91E-05 | 30 | 16 | 12 |
| Temporal_Sup_L | 31 | 4.240839 | 7.73E-05 | -46 | -40 | 16 |
| Olfactory_R | 24 | 3.887782 | 0.000216 | 8 | 28 | -4 |
| Temporal_Mid_L | 31 | 3.7709 | 0.000301 | -56 | -68 | 10 |
| Frontal_Sup_R | 28 | 3.583863 | 0.00051 | 20 | 16 | 60 |
| Temporal_Mid_R | 24 | 3.512822 | 0.000622 | 54 | -26 | -16 |
| Pallidum_L | 10 | 3.465203 | 0.000709 | -12 | 6 | -2 |
| Temporal_Mid_L | 14 | 3.437713 | 0.000765 | -60 | -30 | -8 |
| SupraMarginal_R | 34 | 3.285998 | 0.001157 | 60 | -24 | 30 |
| Precentral_L | 10 | 3.034345 | 0.00226 | -46 | 0 | 20 |

*Note:* All clusters survived the threshold of $p < .005$, with 10 voxel extension, no correction. The names of the anatomical regions in the table, obtained using the AAL toolbox for SPM8, follow the automated anatomical labeling (AAL) template naming convention (Tzourio-Mazoyer et al., 2002). R/L - right/left hemisphere; Ant- anterior; Mid - middle; Sup - superior; Inf - inferior; Orb - orbital; Tri - triangularis. * indicates the voxel cluster that include voxels from the hippocampus.

**Table S3.** Brain regions that showed stronger or weaker activation during *free-viewing*, compared to *fixed-viewing*, of scrambled pictures.

| Anatomical areas | Cluster size | *t* value | *p* value | MNI coordinates | | |
|---|---|---|---|---|---|---|
| | | | | x | y | z |
| **Free-viewing > fixed-viewing** | | | | | | |
| Cuneus_L_R* | 43928 | 10.1982 | 2.53E-12 | 14 | -84 | 20 |
| Frontal_Inf_Oper_R | 2675 | 5.89342 | 5.33E-07 | 46 | 16 | 30 |
| Amygdala_R | 207 | 5.54787 | 1.52E-06 | 34 | -2 | -24 |
| Frontal_Sup_Orb_L | 649 | 5.411448 | 2.31E-06 | -22 | 32 | -14 |
| ParaHippocampal_R | 17 | 4.954296 | 9.21E-06 | 16 | -14 | -24 |
| Temporal_Mid_L | 70 | 4.841744 | 1.29E-05 | -44 | -24 | -12 |
| Cerebelum_9_L | 82 | 4.661586 | 2.22E-05 | -14 | -46 | -48 |
| Frontal_Sup_Medial_R | 92 | 4.526411 | 3.32E-05 | 10 | 62 | 28 |
| Caudate_R | 86 | 4.39434 | 4.92E-05 | 4 | 20 | 4 |
| Frontal_Mid_L | 245 | 4.387761 | 5.02E-05 | -32 | 48 | 32 |
| Insula_R | 128 | 4.250593 | 7.52E-05 | 34 | -18 | 16 |
| Angular_L | 39 | 4.019954 | 0.000147 | -40 | -50 | 24 |
| Frontal_Sup_R | 160 | 4.00789 | 0.000153 | 16 | 40 | 46 |
| Postcentral_R | 120 | 3.79326 | 0.000283 | 66 | -8 | 26 |
| Insula_R | 38 | 3.764184 | 0.000307 | 34 | -4 | 12 |
| Putamen_R | 90 | 3.747869 | 0.000321 | 28 | 16 | 2 |
| Precentral_L | 53 | 3.721141 | 0.000347 | -52 | -4 | 24 |
| Temporal_Pole_Sup_R | 18 | 3.719921 | 0.000348 | 44 | 12 | -24 |
| Caudate_L | 31 | 3.501725 | 0.000641 | -14 | 30 | 2 |
| Thalamus_R | 23 | 3.483542 | 0.000674 | 22 | -10 | 2 |
| Postcentral_L | 51 | 3.426214 | 0.00079 | -48 | -20 | 24 |
| Frontal_Sup_Medial_R | 18 | 3.339045 | 0.001002 | 10 | 54 | 40 |
| Temporal_Sup_R | 40 | 3.321997 | 0.00105 | 46 | -30 | -2 |
| Frontal_Sup_L | 23 | 3.274441 | 0.001194 | -12 | 42 | 50 |
| Temporal_Mid_R | 23 | 3.225212 | 0.001363 | 60 | -42 | 4 |
| Putamen_L | 16 | 3.203533 | 0.001445 | -20 | 4 | 10 |
| Temporal_Sup_R | 20 | 3.187095 | 0.00151 | 64 | -42 | 20 |
| Insula_R | 23 | 3.172441 | 0.00157 | 28 | 18 | -18 |
| Frontal_Mid_L | 17 | 3.159482 | 0.001625 | -26 | 26 | 50 |
| Temporal_Sup_R | 24 | 3.11911 | 0.001808 | 44 | -18 | -8 |
| Temporal_Sup_L | 19 | 3.113383 | 0.001836 | -50 | -28 | 14 |
| **Fixed-viewing > free-viewing** | | | | | | |
| Precuneus_R | 27 | 3.3649 | 0.000934 | 22 | -42 | 16 |

*Note:* All clusters survived the threshold of $p < .005$, with 10 voxel extension, no correction. The names of the anatomical regions in the table, obtained using the AAL toolbox for SPM8, follow the automated anatomical labeling (AAL) template naming convention (Tzourio-Mazoyer et al., 2002). R/L - right/left hemisphere; Ant- anterior; Mid - middle; Sup - superior; Inf - inferior; Orb - orbital; Tri - triangularis. * indicates the voxel cluster that include voxels from the hippocampus.

**Table S4.** Brain regions that showed stronger or weaker (free-viewing > fixed-viewing) viewing effect for the scenes compared to the scrambled pictures.

| Anatomical areas | Cluster size | *t* value | *p* value | MNI coordinates | | |
|---|---|---|---|---|---|---|
| | | | | x | y | z |
| **Scenes > Scrambled** | | | | | | |
| Lingual_R* | 3818 | 9.124077 | 4.41E-11 | 26 | -44 | -8 |
| Lingual_L* | 3334 | 7.963201 | 1.14E-09 | -26 | -48 | -6 |
| Calcarine_L | 202 | 4.317955 | 6.16E-05 | -14 | -50 | 8 |
| Precuneus_R | 242 | 4.250809 | 7.51E-05 | 14 | -52 | 14 |
| **Scenes < Scrambled** | | | | | | |
| Cerebelum_Crus2_L | 178 | 5.365866 | 2.65E-06 | -18 | -74 | -40 |
| Parietal_Inf_L | 693 | 5.09892 | 5.95E-06 | -50 | -48 | 46 |
| Parietal_Sup_R | 1906 | 4.99295 | 8.19E-06 | 48 | -42 | 58 |
| Rolandic_Oper_R | 793 | 4.877378 | 1.16E-05 | 50 | 2 | 18 |
| Supp_Motor_Area_R | 571 | 4.632949 | 2.42E-05 | 16 | 18 | 60 |
| Temporal_Mid_L | 61 | 4.517921 | 3.41E-05 | -60 | -32 | -10 |
| Frontal_Mid_R | 767 | 4.420299 | 4.55E-05 | 44 | 48 | 4 |
| Heschl_R | 513 | 4.367033 | 5.33E-05 | 36 | -26 | 6 |
| Supp_Motor_Area_R | 59 | 4.347381 | 5.65E-05 | 14 | -20 | 56 |
| Supp_Motor_Area_L | 49 | 4.22608 | 8.08E-05 | -12 | 2 | 66 |
| Frontal_Mid_Orb_L | 368 | 4.225111 | 8.1E-05 | -36 | 54 | -4 |
| Cingulum_Mid_R | 232 | 4.01296 | 0.00015 | 20 | -10 | 48 |
| Precentral_L | 129 | 3.99754 | 0.000157 | -26 | -16 | 54 |
| Frontal_Mid_R | 435 | 3.899613 | 0.000208 | 42 | 28 | 40 |
| Cingulum_Mid_L | 51 | 3.88171 | 0.000219 | -14 | -8 | 38 |
| Cerebelum_7b_L | 66 | 3.789446 | 0.000286 | -38 | -52 | -44 |
| Frontal_Sup_L | 103 | 3.786231 | 0.000288 | -12 | 42 | 34 |
| Supp_Motor_Area_R | 27 | 3.668746 | 0.000402 | 6 | -6 | 62 |
| Angular_L | 84 | 3.634419 | 0.000443 | -58 | -58 | 26 |
| Insula_L | 10 | 3.606453 | 0.000479 | -34 | 12 | 14 |
| Temporal_Mid_L | 13 | 3.606255 | 0.000479 | -52 | -44 | -12 |
| Cerebelum_6_L | 24 | 3.558782 | 0.000547 | -18 | -50 | -30 |
| Temporal_Sup_L | 11 | 3.555089 | 0.000553 | -46 | -40 | 14 |
| Frontal_Mid_R | 111 | 3.461617 | 0.000716 | 36 | 40 | 34 |
| Frontal_Mid_L | 55 | 3.439514 | 0.000761 | -24 | 6 | 40 |
| Temporal_Pole_Sup_R | 22 | 3.410617 | 0.000824 | 44 | 10 | -24 |
| Frontal_Sup_L | 55 | 3.379754 | 0.000897 | -14 | 14 | 54 |
| Precentral_L | 61 | 3.357984 | 0.000952 | -48 | 6 | 20 |
| Insula_L | 29 | 3.348258 | 0.000977 | -26 | 18 | 6 |
| Frontal_Inf_Oper_R | 11 | 3.31719 | 0.001064 | 24 | 16 | 30 |
| Frontal_Mid_L | 41 | 3.305507 | 0.001098 | -44 | 30 | 32 |
| Cuneus_R | 34 | 3.303799 | 0.001103 | 8 | -76 | 24 |

| Anatomical areas | Cluster size | *t* value | *p* value | MNI coordinates | | |
|---|---|---|---|---|---|---|
| | | | | x | y | z |
| Precentral_L | 11 | 3.301655 | 0.001109 | -44 | -4 | 38 |
| Frontal_Mid_L | 22 | 3.231787 | 0.001339 | -34 | 20 | 36 |
| Frontal_Sup_R | 14 | 3.17175 | 0.001573 | 20 | 36 | 30 |
| Cuneus_R | 12 | 3.154787 | 0.001645 | 10 | -78 | 36 |
| Cuneus_L | 10 | 3.149628 | 0.001668 | -12 | -76 | 20 |
| Frontal_Mid_L | 13 | 3.080441 | 0.002003 | -38 | 6 | 36 |
| Temporal_Pole_Sup_L | 14 | 3.053262 | 0.002151 | -48 | 8 | -2 |
| Postcentral_R | 19 | 3.004731 | 0.002441 | 62 | -18 | 30 |
| Cingulum_Mid_R | 10 | 2.996816 | 0.002492 | 8 | 36 | 32 |
| Lingual_L | 11 | 2.966526 | 0.002695 | -12 | -60 | -4 |
| Precentral_R | 13 | 2.932666 | 0.002941 | 34 | -24 | 56 |

*Note:* All clusters survived the threshold of $p < .005$, with 10 voxel extension, no correction. The names of the anatomical regions in the table, obtained using the AAL toolbox for SPM8, follow the automated anatomical labeling (AAL) template naming convention (Tzourio-Mazoyer et al., 2002). R/L - right/left hemisphere; Ant- anterior; Mid - middle; Sup - superior; Inf - inferior; Orb - orbital; Tri - triangularis. * indicates the voxel cluster that include voxels from the hippocampus.

**Table S5.** Brain regions for which activations were positively or negatively predicted by the trial-wise number of fixations for scenes in the *free-viewing* condition.

| Anatomical areas | Cluster size | *t* value | *p* value | MNI coordinates | | |
|---|---|---|---|---|---|---|
| | | | | x | y | z |
| **Positive prediction** | | | | | | |
| Calcarine_L | 18321 | 11.01328 | 4.66E-13 | 2 | -88 | -2 |
| Hippocampus_L* | 292 | 5.418666 | 2.45E-06 | -22 | -30 | -2 |
| Parietal_Sup_L | 393 | 4.921602 | 1.09E-05 | -24 | -66 | 52 |
| Caudate_L | 411 | 4.821737 | 1.46E-05 | -22 | -8 | 22 |
| Cerebelum_9_L | 60 | 4.619454 | 2.66E-05 | -16 | -42 | -46 |
| Thalamus_R | 34 | 4.427182 | 4.69E-05 | 22 | -14 | -2 |
| Cerebelum_9_R | 68 | 4.241445 | 8.06E-05 | 16 | -50 | -46 |
| Cerebelum_8_L | 72 | 3.853914 | 0.000246 | -24 | -64 | -44 |
| Amygdala_L | 10 | 3.486204 | 0.000686 | -20 | -4 | -20 |
| Vermis_10 | 60 | 3.409834 | 0.000845 | -4 | -52 | -30 |
| Hippocampus_R* | 33 | 3.391569 | 0.000888 | 38 | -12 | -24 |
| Frontal_Inf_Orb_L | 19 | 3.363371 | 0.000959 | -30 | 32 | -18 |
| Occipital_Mid_L | 13 | 3.198458 | 0.001492 | -22 | -56 | 32 |
| Putamen_L | 11 | 3.159043 | 0.001656 | -22 | 6 | 6 |
| Lingual_R | 12 | 3.055618 | 0.002172 | 10 | -50 | -2 |
| **Negative prediction** | | | | | | |
| Angular_R | 2510 | 6.022786 | 4.02E-07 | 40 | -68 | 48 |
| Precuneus_L | 1638 | 5.752032 | 9.04E-07 | -4 | -56 | 42 |
| Occipital_Mid_L | 689 | 5.709919 | 1.03E-06 | -40 | -76 | 40 |
| Frontal_Sup_R | 4854 | 5.448484 | 2.24E-06 | 18 | 30 | 42 |
| Insula_R | 973 | 5.333827 | 3.17E-06 | 30 | 22 | -6 |
| Temporal_Mid_R | 879 | 5.240514 | 4.19E-06 | 62 | -30 | -12 |
| Frontal_Mid_L | 483 | 5.027965 | 7.9E-06 | -28 | 18 | 54 |
| Insula_L | 409 | 4.779616 | 1.66E-05 | -32 | 18 | -10 |
| Cingulum_Mid_L | 246 | 4.420831 | 4.77E-05 | -8 | 4 | 36 |
| SupraMarginal_L | 384 | 4.272937 | 7.36E-05 | -54 | -24 | 14 |
| Temporal_Mid_L | 57 | 4.119872 | 0.000115 | -66 | -48 | -2 |
| Pallidum_R | 22 | 3.997447 | 0.000163 | 18 | 2 | 0 |
| Temporal_Mid_L | 238 | 3.955889 | 0.000184 | -64 | -24 | -10 |
| Cingulum_Ant_R | 46 | 3.933193 | 0.000196 | 6 | 18 | 20 |
| Frontal_Mid_L | 335 | 3.910862 | 0.000209 | -38 | 52 | 2 |
| Cerebelum_6_L | 10 | 3.854185 | 0.000245 | -24 | -50 | -30 |
| Cingulum_Mid_L | 203 | 3.806512 | 0.000281 | -4 | -22 | 38 |
| Parietal_Inf_L | 56 | 3.721111 | 0.000357 | -50 | -32 | 46 |
| Precentral_R | 20 | 3.604576 | 0.000495 | 24 | -14 | 52 |
| Cerebelum_6_L | 12 | 3.582896 | 0.000525 | -14 | -62 | -26 |

| Anatomical areas | Cluster size | *t* value | *p* value | MNI coordinates | | |
|---|---|---|---|---|---|---|
| | | | | x | y | z |
| Frontal_Inf_Tri_L | 83 | 3.565554 | 0.000551 | -42 | 30 | 24 |
| Vermis_4_5 | 22 | 3.429425 | 0.000801 | 4 | -52 | -14 |
| Precentral_L | 11 | 3.408819 | 0.000847 | -46 | 2 | 20 |
| Postcentral_L | 60 | 3.407134 | 0.000851 | -20 | -30 | 64 |
| Cingulum_Ant_R | 18 | 3.336626 | 0.001031 | 6 | 30 | 8 |
| Postcentral_R | 27 | 3.266626 | 0.001244 | 32 | -40 | 70 |
| Precentral_L | 23 | 3.208074 | 0.001454 | -20 | -18 | 74 |
| Cerebelum_4_5_R | 14 | 3.16168 | 0.001645 | 22 | -48 | -26 |
| Temporal_Sup_L | 16 | 3.095545 | 0.001957 | -44 | 0 | -8 |
| Postcentral_R | 11 | 3.000727 | 0.002505 | 24 | -44 | 60 |
| Parietal_Sup_L | 13 | 2.917289 | 0.003103 | -24 | -46 | 72 |

*Note:* All clusters survived the threshold of $p < .005$, with 10 voxel extension, no correction. The names of the anatomical regions in the table, obtained using the AAL toolbox for SPM8, follow the automated anatomical labeling (AAL) template naming convention (Tzourio-Mazoyer et al., 2002). R/L - right/left hemisphere; Ant- anterior; Mid - middle; Sup - superior; Inf - inferior; Orb - orbital; Tri - triangularis. * indicates the voxel cluster that include voxels from the hippocampus.

**Table S6.** Regions that showed stronger connectivity with the parahippocampal place area (PPA) when scenes were viewed in the *free-* compared to the *fixed-viewing* condition.

| Anatomical areas | Cluster size | *t* value | *p* value | MNI coordinates | | |
|---|---|---|---|---|---|---|
| | | | | x | y | z |
| **Left PPA** | | | | | | |
| Calcarine_R | 11599 | 9.499591 | 1.6E-11 | 8 | -82 | 0 |
| Parietal_Sup_R | 521 | 5.834378 | 6.38E-07 | 26 | -60 | 50 |
| Supp_Motor_Area_L | 285 | 5.720036 | 9.03E-07 | -2 | 8 | 72 |
| Hippocampus_R* | 137 | 5.714706 | 9.18E-07 | 26 | -24 | -4 |
| Hippocampus_L* | 67 | 5.322309 | 3.02E-06 | -20 | -20 | -10 |
| Caudate_R | 175 | 5.31743 | 3.07E-06 | 14 | 28 | -2 |
| Frontal_Inf_Oper_R | 96 | 4.766433 | 1.62E-05 | 40 | 12 | 26 |
| Frontal_Inf_Orb_L | 90 | 4.481842 | 3.79E-05 | -46 | 34 | -12 |
| Frontal_Mid_Orb_R | 17 | 4.396707 | 4.88E-05 | 38 | 38 | -16 |
| Parietal_Sup_L | 199 | 4.286442 | 6.76E-05 | -22 | -64 | 58 |
| Frontal_Sup_Medial_L | 65 | 4.145779 | 0.000102 | -4 | 66 | 26 |
| Precentral_L | 117 | 4.132133 | 0.000106 | -46 | 0 | 58 |
| Cerebelum_10_L | 92 | 4.114556 | 0.000112 | -24 | -38 | -40 |
| Caudate_L | 49 | 4.113555 | 0.000112 | -20 | 12 | 24 |
| Cerebelum_6_L | 27 | 3.794541 | 0.000282 | -30 | -48 | -26 |
| Cerebelum_8_R | 25 | 3.762975 | 0.000308 | 26 | -38 | -48 |
| SupraMarginal_L | 45 | 3.736125 | 0.000332 | -46 | -38 | 30 |
| Putamen_L | 28 | 3.693142 | 0.000375 | -26 | -24 | 4 |
| Occipital_Mid_L | 138 | 3.65078 | 0.000423 | -22 | -92 | 0 |
| Temporal_Inf_L | 13 | 3.604872 | 0.000481 | -44 | -32 | -12 |
| Occipital_Sup_L | 15 | 3.593966 | 0.000496 | -24 | -78 | 42 |
| Frontal_Inf_Tri_R | 53 | 3.585249 | 0.000508 | 48 | 26 | 18 |
| Vermis_3 | 36 | 3.575103 | 0.000523 | 4 | -32 | -4 |
| Temporal_Inf_R | 24 | 3.561748 | 0.000543 | 44 | -12 | -26 |
| Fusiform_R | 10 | 3.461729 | 0.000716 | 20 | 14 | -44 |
| Frontal_Inf_Orb_L | 14 | 3.450275 | 0.000739 | -34 | 36 | -10 |
| Caudate_R | 38 | 3.320849 | 0.001053 | 12 | -6 | 24 |
| Thalamus_L | 14 | 3.317949 | 0.001061 | -2 | -14 | 16 |
| Frontal_Inf_Tri_R | 12 | 3.291466 | 0.00114 | 56 | 38 | 16 |
| Caudate_L | 20 | 3.232363 | 0.001337 | -12 | 22 | 12 |
| Precentral_R | 10 | 3.214593 | 0.001402 | 56 | 12 | 40 |
| Precuneus_R | 10 | 3.19248 | 0.001488 | 10 | -66 | 68 |
| Cerebelum_9_R | 10 | 3.134873 | 0.001734 | 14 | -44 | -44 |
| **Right PPA** | | | | | | |
| Occipital_Inf_R | 14985 | 11.201 | 2.01E-13 | 40 | -80 | -4 |
| Hippocampus_L* | 124 | 6.395288 | 1.17E-07 | -22 | -24 | -6 |

| Anatomical areas | Cluster size | *t* value | *p* value | MNI coordinates | | |
|---|---|---|---|---|---|---|
| | | | | x | y | z |
| Frontal_Mid_Orb_L | 73 | 6.105368 | 2.81E-07 | -34 | 36 | -14 |
| Thalamus_R* | 130 | 5.618295 | 1.23E-06 | 24 | -22 | -4 |
| Supp_Motor_Area_R | 303 | 5.472495 | 1.92E-06 | 4 | 12 | 66 |
| Frontal_Inf_Tri_R | 156 | 5.258107 | 3.67E-06 | 54 | 32 | 0 |
| Caudate_R | 157 | 5.157291 | 4.98E-06 | 24 | -30 | 24 |
| Cerebelum_7b_L | 41 | 4.814644 | 1.4E-05 | -6 | -76 | -40 |
| Precentral_L | 101 | 4.73292 | 1.79E-05 | -48 | -4 | 50 |
| Cingulum_Ant_R | 263 | 4.66837 | 2.18E-05 | 18 | 34 | 6 |
| Precentral_R | 25 | 4.481076 | 3.8E-05 | 38 | -14 | 38 |
| Lingual_R | 60 | 4.404961 | 4.77E-05 | 6 | -30 | -4 |
| Caudate_L | 66 | 4.392404 | 4.95E-05 | -18 | 32 | 0 |
| Frontal_Inf_Tri_L | 215 | 4.298525 | 6.53E-05 | -54 | 34 | 2 |
| Frontal_Inf_Tri_R | 152 | 4.132302 | 0.000106 | 38 | 18 | 20 |
| Frontal_Inf_Orb_R | 51 | 4.024198 | 0.000146 | 34 | 34 | -22 |
| Cerebelum_10_L | 28 | 3.851404 | 0.000239 | -18 | -40 | -44 |
| Temporal_Pole_Sup_R | 35 | 3.740122 | 0.000329 | 34 | 22 | -26 |
| Frontal_Sup_Medial_L | 36 | 3.688224 | 0.000381 | -2 | 66 | 18 |
| Precentral_L | 13 | 3.680204 | 0.000389 | -26 | -12 | 48 |
| Parietal_Sup_L | 131 | 3.643467 | 0.000432 | -20 | -52 | 52 |
| Rolandic_Oper_L | 14 | 3.632471 | 0.000445 | -48 | -16 | 22 |
| Temporal_Pole_Sup_L | 14 | 3.59922 | 0.000489 | -46 | 24 | -26 |
| Rolandic_Oper_R | 11 | 3.448575 | 0.000743 | 48 | -6 | 8 |
| Frontal_Sup_L | 18 | 3.365325 | 0.000933 | -28 | -2 | 68 |
| Occipital_Mid_L | 14 | 3.291732 | 0.001139 | -48 | -76 | 8 |
| Caudate_L | 10 | 3.247858 | 0.001283 | 0 | 12 | 4 |
| Precentral_R | 14 | 3.207404 | 0.00143 | 24 | -30 | 68 |
| Frontal_Mid_R | 12 | 3.189278 | 0.001501 | 46 | 0 | 58 |
| Precentral_R | 17 | 3.185396 | 0.001516 | 52 | 2 | 50 |
| Supp_Motor_Area_L | 11 | 3.178357 | 0.001545 | -10 | -6 | 64 |
| Frontal_Sup_Medial_L | 16 | 3.165672 | 0.001598 | -2 | 52 | 44 |
| Temporal_Inf_L | 13 | 3.150594 | 0.001664 | -34 | 4 | -36 |

*Note:* All clusters survived the threshold of $p < .005$, with 10 voxel extension, no correction. The names of the anatomical regions in the table, obtained using the AAL toolbox for SPM8, follow the automated anatomical labeling (AAL) template naming convention (Tzourio-Mazoyer et al., 2002). R/L - right/left hemisphere. * indicates the voxel cluster that include voxels from the hippocampus.

**Table S7.** Brain regions that showed subsequent memory effects for scenes in the *free-viewing* condition.

| Anatomical areas | Cluster size | *t* value | *p* value | MNI coordinates | | |
|---|---|---|---|---|---|---|
| | | | | x | y | z |
| **Remembered > forgotten** | | | | | | |
| Occipital_Sup_R* | 5385 | 7.44754 | 6.1E-09 | 26 | -68 | 38 |
| Precentral_R | 742 | 6.510548 | 9.43E-08 | 46 | 4 | 32 |
| Occipital_Mid_L* | 4352 | 6.189985 | 2.44E-07 | -34 | -88 | 22 |
| Cerebelum_9_L | 82 | 5.200029 | 4.72E-06 | -14 | -46 | -48 |
| Thalamus_R | 64 | 5.022265 | 8.04E-06 | 18 | -30 | 2 |
| Frontal_Inf_Oper_L | 262 | 4.41166 | 4.9E-05 | -44 | 8 | 28 |
| Caudate_R | 33 | 4.158551 | 0.000103 | 26 | 8 | 18 |
| Thalamus_R | 44 | 4.098952 | 0.000122 | 2 | -14 | 4 |
| Frontal_Inf_Orb_L | 26 | 3.866363 | 0.000237 | -32 | 36 | -16 |
| Precuneus_L | 40 | 3.813852 | 0.000275 | -16 | -50 | 10 |
| Insula_R | 25 | 3.652804 | 0.000432 | 46 | 6 | 6 |
| Insula_L | 35 | 3.652133 | 0.000433 | -34 | -2 | -14 |
| Putamen_L | 14 | 3.492128 | 0.000675 | -22 | -6 | 12 |
| Cerebelum_9_L | 53 | 3.384715 | 0.000905 | -2 | -52 | -34 |
| Frontal_Inf_Orb_R | 36 | 3.316984 | 0.001087 | 28 | 32 | -12 |
| Caudate_L | 34 | 3.251357 | 0.001296 | -8 | 8 | 18 |
| Supp_Motor_Area_R | 34 | 3.236993 | 0.001347 | 8 | 12 | 54 |
| Putamen_L | 12 | 3.159378 | 0.001655 | -20 | 6 | 6 |
| Thalamus_R | 23 | 3.049127 | 0.002209 | 10 | -20 | 6 |
| **Forgotten > remembered** | | | | | | |
| Cingulum_Post_R | 6388 | 7.447096 | 6.11E-09 | 2 | -48 | 30 |
| Angular_R | 3762 | 6.425117 | 1.21E-07 | 52 | -52 | 36 |
| Cingulum_Ant_R | 9569 | 6.187421 | 2.46E-07 | 4 | 44 | 26 |
| Insula_R | 253 | 6.055739 | 3.65E-07 | 32 | 20 | -10 |
| Angular_L | 4483 | 5.897652 | 5.85E-07 | -46 | -56 | 38 |
| Precentral_L | 152 | 4.863593 | 1.29E-05 | -26 | -14 | 66 |
| Temporal_Pole_Sup_R | 32 | 4.170016 | 9.92E-05 | 56 | 10 | -12 |
| Postcentral_R | 71 | 4.159064 | 0.000102 | 20 | -44 | 64 |
| Cerebelum_Crus2_L | 66 | 4.106555 | 0.000119 | -24 | -86 | -34 |
| Insula_L | 89 | 4.104821 | 0.00012 | -32 | 16 | -10 |
| Cingulum_Ant_L | 50 | 4.059104 | 0.000137 | -4 | 18 | 20 |
| Cingulum_Post_R | 80 | 3.978703 | 0.000172 | 10 | -36 | 12 |
| Lingual_L | 80 | 3.96981 | 0.000177 | -12 | -56 | -4 |
| Lingual_R | 119 | 3.684425 | 0.000396 | 12 | -58 | -6 |
| Supp_Motor_Area_R | 11 | 3.478273 | 0.000701 | 12 | -22 | 56 |

| Anatomical areas | Cluster size | *t* value | *p* value | MNI coordinates | | |
|---|---|---|---|---|---|---|
| | | | | x | y | z |
| Frontal_Inf_Tri_R | 48 | 3.472286 | 0.000712 | 48 | 20 | 2 |
| Precentral_L | 10 | 3.47227 | 0.000712 | -34 | -4 | 38 |
| Frontal_Mid_L | 37 | 3.402615 | 0.000862 | -38 | 38 | 22 |
| Temporal_Mid_R | 11 | 3.331515 | 0.001045 | 52 | -4 | -28 |
| Postcentral_L | 35 | 3.263992 | 0.001253 | -48 | -24 | 56 |
| Temporal_Pole_Mid_R | 17 | 3.206362 | 0.001461 | 46 | 16 | -30 |

*Note:* All clusters survived the threshold of $p < .005$, with 10 voxel extension, no correction. The names of the anatomical regions in the table, obtained using the AAL toolbox for SPM8, follow the automated anatomical labeling (AAL) template naming convention (Tzourio-Mazoyer et al., 2002). R/L - right/left hemisphere; Ant- anterior; Mid - middle; Sup - superior; Inf - inferior; Orb - orbital; Tri - triangularis. * indicates the voxel cluster that include voxels from the hippocampus.

**Supplementary Figure S1:** ROI masks used in the present study

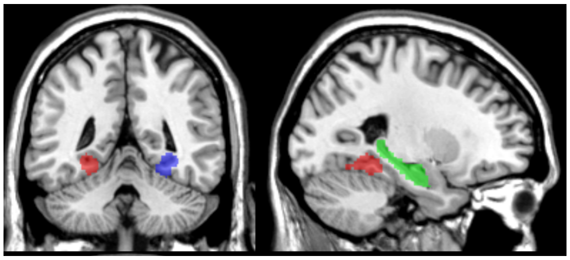

Green = hippocampus; red = left PPA; blue = right PPA. PPA = parahippocampal place area.
